# The branching code: a model of actin-driven dendrite arborisation

**DOI:** 10.1101/2020.10.01.322750

**Authors:** Tomke Stürner, André Ferreira Castro, Maren Philipps, Hermann Cuntz, Gaia Tavosanis

**Author notes:** Senior authors (Hermann Cuntz, Gaia Tavosanis). Correspondence (G.T.), (H.C.).

## Abstract

Dendrites display a striking variety of neuronal type-specific morphologies, but the mechanisms and principles underlying such diversity remain elusive. A major player in defining the morphology of dendrites is the neuronal cytoskeleton, including evolutionarily conserved actin-modulatory proteins (AMPs). Still, we lack a clear understanding of how AMPs might support developmental phenomena such as neuron-type specific dendrite dynamics. To address precisely this level of *in vivo* specificity, we concentrated on a defined neuronal type, the class III dendritic arborisation (c3da) neuron of *Drosophila* larvae, displaying actin-enriched short terminal branchlets (STBs). Computational modelling reveals that the main branches of c3da neurons follow a general growth model based on optimal wiring, but the STBs do not. Instead, model STBs are defined by a short reach and a high affinity to grow towards the main branches. We thus concentrated on c3da STBs and developed new methods to quantitatively describe dendrite morphology and dynamics based on *in vivo* time-lapse imaging of mutants lacking individual AMPs. In this way, we extrapolated the role of these AMPs in defining STB properties. We propose that dendrite diversity is supported by the combination of a common step, refined by a neuron type-specific second level. For c3da neurons, we present a molecular model of how the combined action of multiple AMPs *in vivo* define the properties of these second level specialisations, the STBs.

**In brief:** A quantitative morphological dissection of the concerted actin-modulatory protein actions provides a model of dendrite branchlet outgrowth.

**Highlights:**

- Actin organisation in small terminal branchlets of *Drosophila* class III dendritic arborisation neurons
- Six actin-modulatory proteins individually control the characteristic morphology and dynamics of branchlets
- Quantitative tools for dendrite morphology and branch dynamics enable a comparative analysis
- A two-step computational growth model reproduces c3da dendrite morphology

## Introduction

Regulated outgrowth and branching are essential to establish neuronal dendrites optimised to perceive and appropriately process specific inputs (Jan and Jan, 2010). This functional requirement defines clear structural constraints. Features of dendrite morphology are thus tightly correlated to neuronal function and are distinctive enough to enable a first level of neuron type classification (MacNeil and Masland, 1998). The process of how neuronal type-specific dendrite morphology can be achieved is a key question for elucidating the development of the nervous system.

Core aspects of dendrite morphology are defined by transcription factor codes determined during development and that impart neuronal identity (Santiago and Bashaw, 2014; Dong et al., 2015; Parrish et al., 2007; Ziegler et al., 2017). In addition to intrinsic factors, signals derived from a neuron’s environment, including those that support the establishment of functional connections, contribute to refining dendritic structure (Corty et al., 2009; Valnegri et al., 2015; Dong et al., 2015). These multiple layers of regulation converge on the control of the cellular cytoskeleton, which ultimately defines the structural and dynamical properties of cells (Konietzny et al., 2017; Coles and Bradke, 2015). The ensemble of numerous AMPs, in particular, drives the dynamics that lead to dendritic tree establishment (Lanoue and Cooper, 2019). Most key AMPs are highly conserved across species and their biochemical properties have been carefully analysed *in vitro* (Mullins et al., 1998; Pruyne et al., 2002; Breitsprecher et al., 2008; Kovar et al., 2006; Smith et al., 2013) and in cultured cells (Damiano-Guercio et al., 2020; Suraneni et al., 2012; Wu et al., 2012; Koestler et al., 2013). The collective activity of various AMPs describe different protrusion types during cell migration (Schaks et al., 2019). However, our understanding of how AMPs cooperate in space and time to form specialised dendritic morphologies during animal development is still highly speculative (Konietzny et al., 2017).

The dendritic arborisation (da) neurons of *Drosophila melanogaster* represent a fruitful system for studying the complex role of actin and AMPs in dendrite morphogenesis *in vivo* (Corty et al., 2009). Four morphologically and functionally distinct classes of da neurons (c1da–c4da) extend their planar dendrites underneath the larval transparent cuticle facilitating live imaging of their differentiation. In particular, the dynamics of actin organisation can be studied *in vivo* in these neurons using genetically encoded fluorescent fusion proteins that associate with actin filaments (Kiehart et al., 2000; Hatan et al., 2011; Haralalka et al., 2014; Nithianandam and Chien, 2018). These tools have allowed the visualisation of localised dynamic actin accumulation preceding new branch formation (Andersen et al., 2005; Stürner et al., 2019).

In combination with the imaging efforts, genetic studies have involved multiple cytoskeletal regulators in the establishment of da dendrites *in vivo*. The actin severing and depolymerising protein Twinstar / cofilin regulates actin at dendrite branching sites in the c4da neurons and supports branch formation in all da classes (Nithianandam and Chien, 2018). The actin nucleator complex Arp2/3 transiently localises at branching sites where it forms branched actin to initiate branchlet formation in all da neuron classes (Stürner et al., 2019). The actin barbed end binding protein Ena / VASP promotes lateral branching of all da neuron classes (Gao et al., 1999; Dimitrova et al., 2008). While these AMPs seem to cover a general function in branch formation, others can be neuron type-specific. A striking example is afforded by the actin bundling protein Singed / fascin, which localises exclusively within the terminal branchlets of c3da neurons and is required only in this distinctive type of branchlet (Nagel et al., 2012). In addition, the actin nucleation factor Spire is differentially regulated in c1da and c4da neurons (Ferreira et al., 2014). The latter studies indicate that individual subsets of branches even within a neuron contain specific AMPs defining their morphological and dynamic properties. Furthermore, they seem to suggest that a core, general program supporting dendrite establishment exists, but that this general program needs to be associated with a neuron type-specific secondary program to define the morphology of specific neuron types.

To understand how specific AMPs work in concert to control dendrite branchlet properties and thus regulate dendrite morphology we focused on one type of dendritic branch in one class of da neurons. The c3da neurons of *Drosophila* larvae respond to gentle touch (Tsubouchi et al., 2012; Yan et al., 2013) and noxious cold (Turner et al., 2016). They display long primary dendrite branches decorated with characteristic short and dynamic terminal branchlets (STBs) (Grueber et al., 2002; Andersen et al., 2005; Nagel et al., 2012) that are required for gentle touch responses (Tsubouchi et al., 2012; Yan et al., 2013). The c3da STBs are highly enriched in actin, making them an ideal model system to study actin-dependent branching dynamics *in vivo*.

Early studies exploring the functional role of AMPs in dendrite elaboration often relied on single static features or morphometrics, such as the number of branches for a given Strahler order or Sholl analysis (Ferreira et al., 2014; Vormberg et al., 2017; Bird and Cuntz, 2019; Kanaoka et al., 2019). While such approaches can reveal the involvement of AMPs, they might fall short of pointing to the specific role of individual AMPs. Recently, morphological modelling has proven to be an important method to probe our understanding of dendritic morphology that can additionally point to a new mechanistic insight of development (Cuntz, 2016; Poirazi and Papoutsi, 2020). In such a morphological modelling approach, synthetic morphologies are built from a set of assumptions made about branching statistics (e.g. Koene et al., 2009; Ascoli et al., 2001), wiring considerations (e.g. Cuntz et al., 2007, 2008; Budd et al., 2010; Cuntz et al., 2010), their underlying growth rules (e.g. Sugimura et al., 2007; Memelli et al., 2013; Torben-Nielsen and Schutter, 2014) or even the computation that a given neuron is thought to implement (Torben-Nielsen and Stiefel, 2010). In selected cases, morphological modelling of specific dendrites has elucidated the logic underlying their structural plasticity during maturation (Beining et al., 2017) or specific manipulations (Sugimura et al., 2007; Nanda et al., 2018a,b, 2019; Yalgin et al., 2015).

Specifically, a novel growth model was designed recently that has been fitted to the details of c4da dendrite growth during larval development (Baltruschat et al., 2020). This model is particularly interesting since it both reproduces the branching behaviour of these cells and satisfies the more mathematical aspects of dendrite morphological modelling derived from space filling and optimal wiring criteria. Thereby, the model links a phenomenological description of mature dendrite morphology with the biological processes that shape their growth dynamics and lead to the mature dendrites in the final stages of larval development. Moreover, the iterations of growth described by the model translate directly to the rough description of dendrites in other cell types including three dimensional dendrites in mammalian cortex such as dentate gyrus granule cells and cortical pyramidal cells in various layers (Baltruschat et al., 2020). With a more detailed modelling approach this general growth model derived from c4da neurons has also been applied to understand the dendritic computations performed by c1da neurons in the fly larva (Castro et al., 2020). We thus took advantage of the possibility of linking dynamics of dendrite growth with a more formal and mathematical understanding of dendrite morphology afforded by the c4da model to dissect the dynamic growth process of c3da neuron dendrites.

In this study, we imaged the morphology and dynamics of c3da dendrites *in vivo* in wild-type animals or in mutants of four AMP genes important for defining c3da neuron morphology. Utilising improved quantitative analysis we could assign discrete roles to each of these factors. Additionally, we revealed novel roles for two additional AMPs, Spire and Capuccino (Capu). We further produced a two-step growth model that can accurately replicate the characteristic wild-type c3da neuron dendrite morphology and applied it to each of the mutants. We thus put forward a comprehensive model of actin-regulated control of c3da STB dynamics in the context of a two-step computational model of c3da neuron morphology.

## Results

### A two-step model is necessary to describe the c3da neuron morphology

We used computational modelling as a first step towards understanding the characteristic morphology of c3da neurons and which growth rules could apply to their dendrite morphology. C3da neurons tile covering 70% of the body wall and scale during the larval growth phase, similarly to c4da neurons (Grueber et al., 2002; Parrish et al., 2009). We therefore first used a model that we recently developed for c4da neuron dendrites based on their ability to innervate their target area in a space-filling manner (Baltruschat et al., 2020). This space-filling growth model that accurately reproduces the development of c4da dendrites is based on previous models that satisfy optimal wiring constraints by balancing costs for total dendritic length and signal conduction times (Cuntz et al., 2007, 2008, 2010, 2012). It utilises simple parameters such as the target spanning area, a value for stochasticity of innervation (*k*) and a factor *(bf*) representing the balance between total dendrite length and path length to the soma as defined in (Cuntz et al., 2007, 2010, see **STAR**\***Methods**). The growth model replicates the general features of dendrite morphology in a wide variety of neuronal cell types, including Purkinje cells, hippocampal granule and pyramidal cells, as well as cortical pyramidal cells (Baltruschat et al., 2020). Thus, it seems to well represent core general properties of dendrite morphology establishment and we refer to it as the general growth model throughout this work.

To model their morphology, we first imaged control ldaB c3da neurons of the abdominal segment A5 of early third instar larvae (L3) *in vivo* and traced them in 3D in the *TREES toolbox* (www.treestoolbox.org; Cuntz et al., 2010). Similarly as performed for c4da neurons (see details in Baltruschat et al., 2020), we let the general growth model described above innervate the spanning area covered by the reconstructed c3da neurons (**Figure 1A**, grey shade, see **STAR**\***Methods**). We first focused on the main branches of the c3da neuron by removing all terminal branches and then recursively all terminal branches shorter than 10μm until none were left (**Figure 1A**, left). We found fitting parameters for the model with a *bf* of 0.1, a low *k* of 0.15, a radius reach of 100*μm* (see **STAR**\***Methods** for more details). These parameters did not differ much from the model directly simulating c4da dendrite growth (Baltruschat et al., 2020). To obtain this, the simulated growth process was stopped when the number of branches reached the number of main branches in the corresponding real dendrite (**Figure 1A**, middle). Total length and overall shape were then similar to the real counterparts (**Figure 1A**, right). However, when resuming growth in the model, the new branches filled the available space failing to reproduce the characteristic STBs observed in c3da dendrites (compare **Figure 1B** left and middle morphologies as well as the corresponding Sholl intersections on the right).

**Fig 1.**
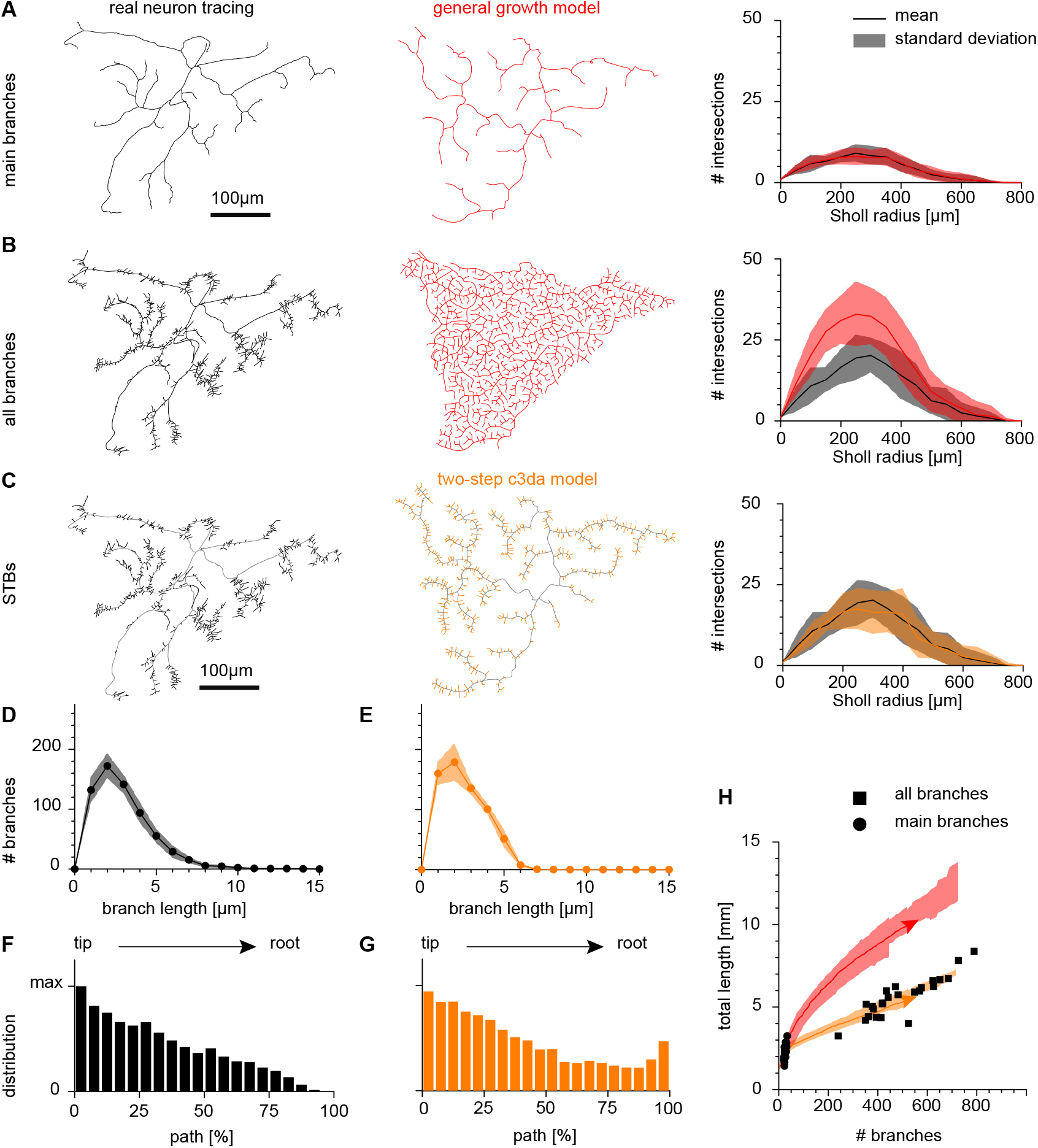
A two-step c3da model. **A**, **B**, **C**, Tracings of a wild type c3da neuron (black) with spanning area (shaded in grey) and synthetic dendritic trees (red or orange) focusing on the main branches (**A**), all branches (**B**) or the STBs (**C**). Right hand Sholl analysis panels show the number of intersections of the dendritic trees with increasing Sholl radii around the soma in *μm.* Shaded area shows standard deviation. Solid lines show the mean Sholl intersections. **A**, **B**, The synthetic dendritic trees in red were generated with the general growth model (Baltruschat et al., 2020), but the growth was interrupted either when the number of main branches in **A** was reached or interrupted when the total number of branches in **B** was reached. **C**, A second modelling step of the synthetic dendritic tree in orange allows STBs with a defined total length to develop in a close range to the main branch with a given distribution along the main branches. **D**,**E**, The number of STBs in the real neuron tracings (**D**) and the synthetic trees obtained with the two-step model (**E**) plotted against their length in μm. **F**,**G**, The number of STBs at positions along the main branches, from tip to root (depicted as a percentile of the path length). **H**, Number of branches vs. total length for main branches (black dots) and complete trees (black squares). Trajectories with standard deviation are shown for the general growth model (shaded red area) and the two-step c3da model (shaded orange area). Solid arrows show examples in **B** and **C**, respectively. Scale bar is 100*μm* (see **STAR**\***Methods** for details and **Table 2** for genotypes).

Taken together, the general growth model by Baltruschat et al. (2020) that successfully reproduces the dendrite morphology of space-filling neurons is not sufficient to describe c3da neurons because of the number, shape and distribution of their characteristic STBs. In line with these findings, c3da neurons were previously singled out for the irregular distribution of their branches (Anton-Sanchez et al., 2018). Computational modelling of these neurons thus seems to require more restrictions than optimal wiring and space filling and needs to include the distinction between main branches and STBs. After preserving the main branches in accordance with the space-filling growth model as demonstrated in **Figure 1A, B**, we added STBs in a second growth phase. This second phase was intentionally kept as similar as possible to the general growth model to be able to identify the distinct differences between STBs and the main branches of c3da dendrites. This second step in the growth model required different parameters *bf =* 0.65 and *k* = 0.5 and a much closer reach around the main branches that correlated with the distance to the root. Most importantly, STBs grew with a specific affinity towards the main branches rather than to the root of the entire dendrite making this growth rule markedly distinct from other growth rules described previously (see details in **STAR**\***Methods**).

Based on the dendrite total length this two-step model derived a branch length distribution of STBs along the main branch that was almost indistinguishable from that of the real counterparts as demonstrated with Sholl intersection diagrams (**Figure 1C**). The addition of STBs as a second step led to the replication of the characteristic branch length distribution of STBs (**Figure 1D, E**). The new synthetic dendritic trees visually resembled the wild type, displaying a similar distribution probability of the STBs along the main branches (**Figure 1F, G**, see **STAR**\***Methods** for details). The c3da wild-type trees aligned with the growth trajectories obtained using this two-step c3da model with respect to their number of branches and total length (**Figure 1H**), lying well off the trajectories predicted by the general growth model.

### Actin organisation in the short terminal branchlets of c3da neurons

The model singled out the STBs as a second, neuron-type specific level of dendrite elaboration of c3da neurons. STBs of c3da neurons are actin- and Singed / fascin-enriched straight branchlets which dynamically extend and retract throughout larval stages (Nagel et al., 2012). To understand how these branches are formed and how their dynamics are coordinated by AMPs, we first investigated the organisation and dynamics of the actin cytoskeleton *in vivo*. To define the orientation of the actin filaments and their dynamic properties we performed a fluorescence recovery after photobleaching (FRAP) analysis of green fluorescent protein (GFP)-labelled actin in the STBs of lateral c3da neurons (**Figure 2A,B**). For an internal reference, we also expressed a fluorescent, membrane-targeted chimeric protein highlighting the dendritic branchlet.

**Fig 2.**
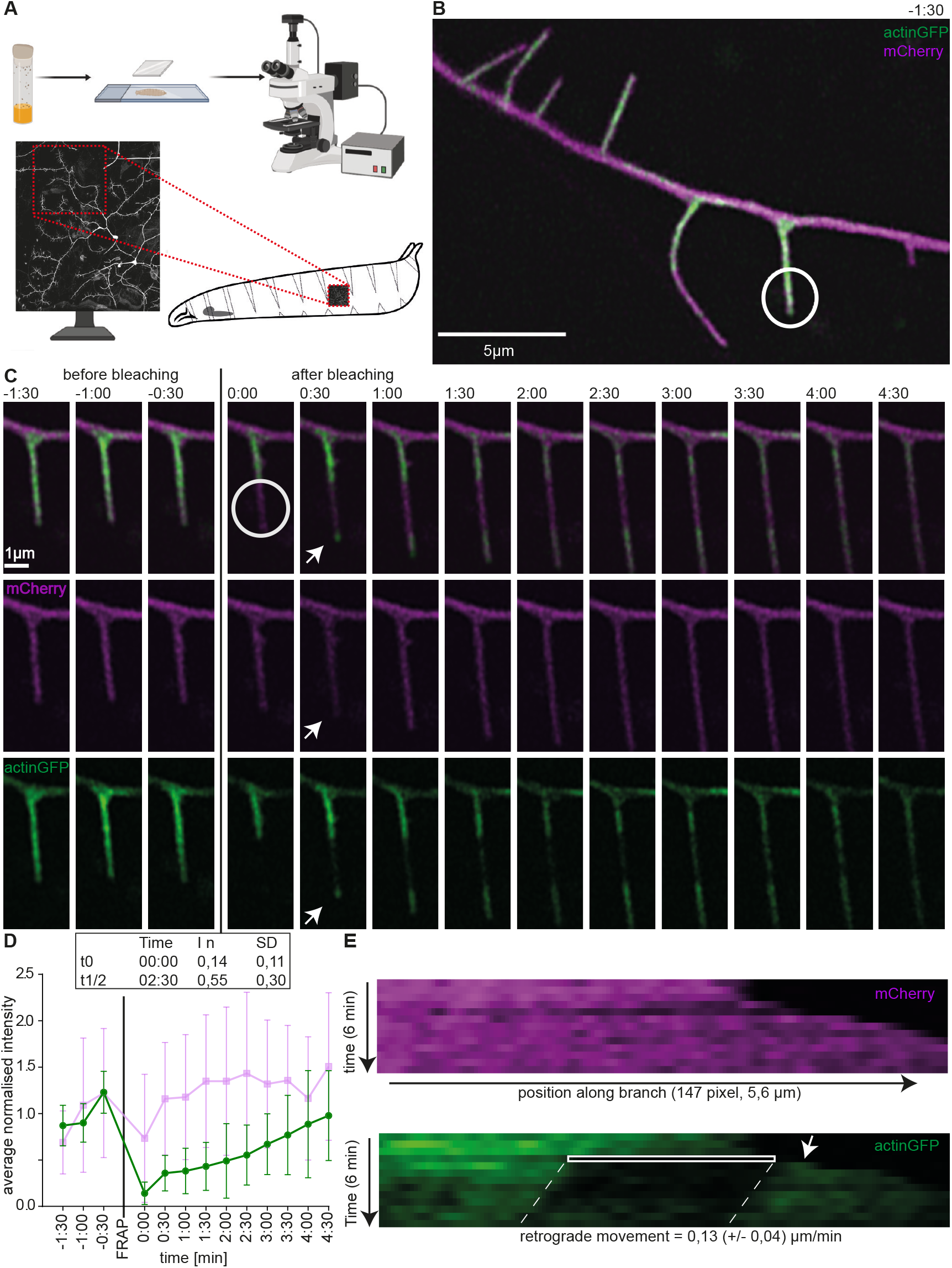
FRAP analysis of actin in c3da neuronal terminal branches. **A**, Illustration of the setup for time-lapse imaging of ldaB c3da neurons. Terminal branches for time-lapse imaging were chosen in a defined dendrite quadrant (red square). Image created with BioRender.com. **B**, Representative overview image of a c3da dendritic branch 1*min* before bleaching. *UASmCD8Cherry* – Magenta and *UASp – GFP.Act5C* – Green. The white circle indicates the photobleached area at time point 0:00 of the time-lapse series. **C**, Time-lapse images of the same STB (from **B**) are shown every 30*sec* over a 6*min* interval. The white circle indicates the photobleached area at time point 0:00. The white arrow points to the bright GFP signal at the growing branchlet tip after photobleaching. **D**, Average normalised Actin-GFP and Membrane-mCherry fluorescence intensity in the bleached area of 8 time series. **E**, A representative kymograph of the same dendritic branchlet over time and space. The bleached area is highlighted with a white rectangle and dashed white lines indicate the retrograde movement of filamentous actin in this area, *r*. The white arrow points to the bright actin-GFP signal recovery after photobleaching. *n* = 8 neurons from individual larvae (see **STAR**\***Methods** for details and **Table 2** for genotype).

While the membrane-targeted chimera signal was almost unaffected, the actin::GFP signal dropped to 0.14*In* after photobleaching (**Figure 2C,D**, see **STAR**\***Methods** for details). After bleaching only the tips of elongating dendritic branchlets (white circle in **Figure 2B**) we examined where new actin monomers are added to the actin filaments (**Figure 2A,B,C**). Merely 30*sec* after photobleaching, the tip of elongating dendritic branchlets displayed a sharp recovery of actin::GFP signal at the distal end of the bleached area (**Figure 2C**, arrow). Thus branchlet elongation correlated with actin filament elongation at the extending distal tip of the branchlet. Thus, c3da STBs contain mostly actin filaments with their fast-growing ends pointing distally.

We tracked the length and fluorescence intensity of the branchlet over time and measured the actin::GFP signal within the bleached area (see `analysisFRAP_macro.ijm`, **Figure 2C,D,E**), revealing the velocity of actin turnover (half-time recovery 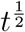) and the speed of actin treadmilling (retrograde movement *r,* **Figure 2E,E**) (Lai et al., 2008). The average half-time of recovery of actin::GFP in the bleached area was 2.5*min* after photobleaching (**Figure 2D**; 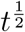) and full actin recovery in c3da terminal branchlets was around 5*min* (**Figure 2D**). Within the bleached area the Actin::GFP signal recovered evenly, suggesting that the bundle harboured interspersed actin filaments of different length (**Figure 2C,E**).

A kymograph of actin GFP fluorescence visualised the treadmilling of actin within the growing branchlet (**Figure 2E**). The retrograde movement velocity of the bleached area in the present study was 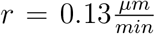. Taken together and given also the known enrichment of Singed / fascin (Nagel et al., 2012), c3da STBs apparently contain mainly uniparallel actin bundles oriented with the majority of fast growing ends pointing distally and displaying slow actin kinetics.

### Analysis of six AMPs that regulate dendrite branch number in c3da neurons

To identify the molecular regulation of actin in the c3da neuron dendrites, we performed literature searches and a targeted screen of actin nucleators (Stürner et al., 2019), elongators, bundling and depolymerisation factors. We concentrated our analysis on mutants of six AMPs and imaged their c3da neurons *in vivo* at the early third instar larva stage (**Figure 3A**, see **Table 1** for fly strains). To extract a deep quantitative phenotypic description of their dendrite morphology, we traced and analysed the c3da neuron images in the *TREES toolbox* (www.treestoolbox.org; Cuntz et al., 2010).

**Fig 3.**
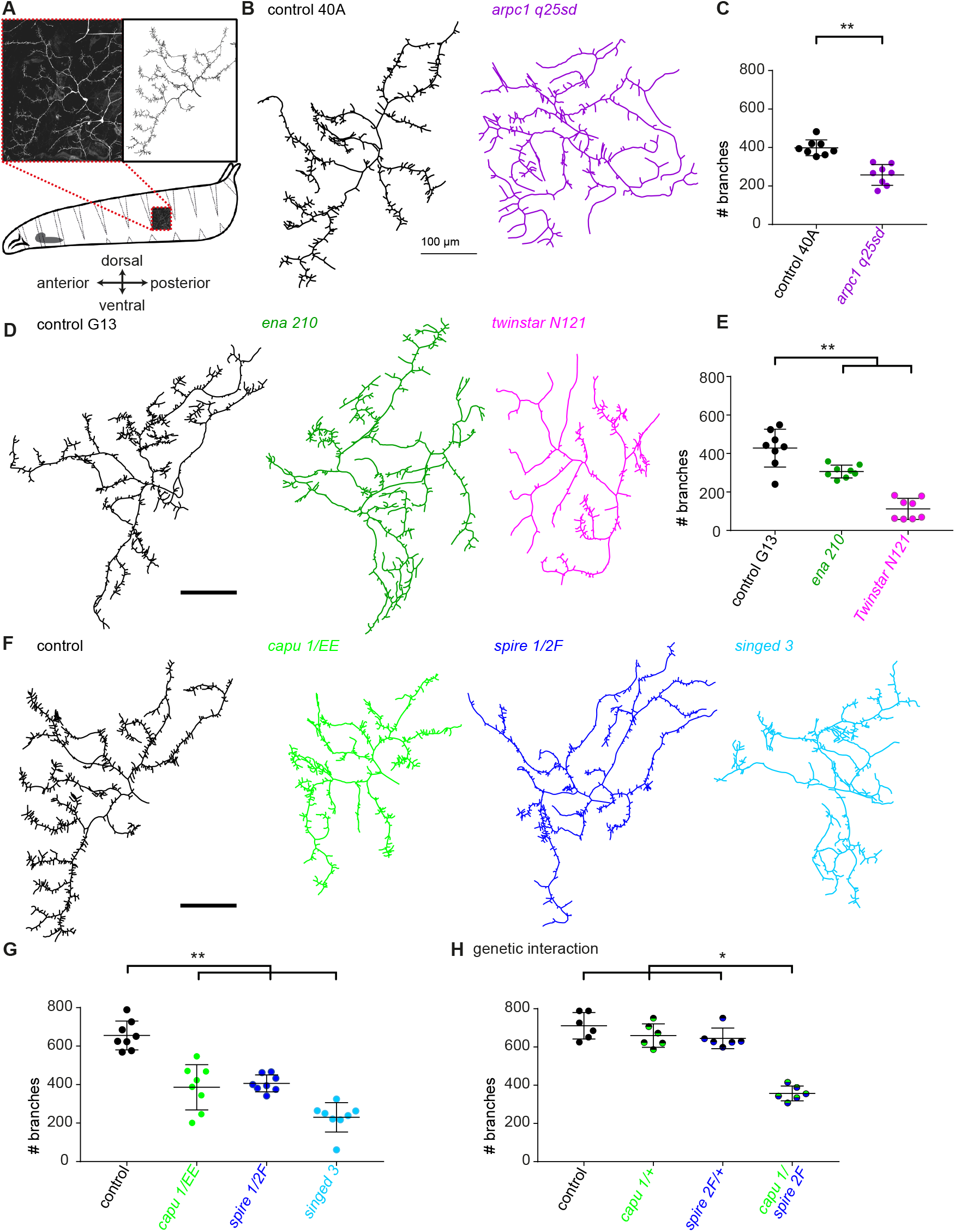
Actin modulatory proteins involved in terminal branch formation of c3da neurons. **A**, Illustration of c3da neuron imaging and tracing reconstructed in the *TREES toolbox.* All tracings are shown in the orientation as shown in this scheme. **B**, Representative tracing of MARCM clones of control and *arpc*1^*q*25*sd*^ mutants. **D**, Representative tracing of MARCM clones of control, ena^210^ mutants and *tsr*^*N*121^ mutants. **F**, Representative tracing of control, *capu*^1^ /*capu^EE^*, *spire*^1^ /*spire*^2*F*^ and *singed*^3^ mutants. **C**, **E**, **G**, Quantification of total branch number of the different groups with controls. **H**, Quantification of total branch number in heterozygous mutants of *spire*^2*F*^/+, *capu*^1^ /+, or *capu*^1^ /*spire*^2*F*^ transheterozygous mutants. (* is *p* < 0.05, ** is *p* < 0.01 and *** is *p* < 0.001). Scale bar is 100*μm*. *n* = 8 neurons from individual larva per genotype (see **Table 2** for genotypes).

**Table 1.** Reagent and Resource.

| REAGENT or RESOURCE | SOURCE | IDENTIFIER |
| --- | --- | --- |
| Experimental Models: Organisms/Strains |  |  |
| *D. melanogaster*: *P{ry[+t7.2]=hsFLP}12, y[1] w[\*]; Arpc1[Q25sd] P{ry[+t7.2]=neoFRT}40A/CyO* | Bloomington Drosophila Stock Center | BDSC: 9137 |
| *D. melanogaster*: *spir[1] cn[1] bw[1]/CyO, l(2)DTS513[1]* | Bloomington Drosophila Stock Center | BDSC: 5113 |
| *D. melanogaster*: *b[1] pr[1] spir[2F] cn[1]/CyO* | Bloomington Drosophila Stock Center | BDSC: 8723 |
| *D. melanogaster*: *M{UAS-spir. ORF. 3xHA}ZH-86Fb* | FlyORF | F001174 |
| *D. melanogaster*: *capu[1] cn[1] bw[1]/CyO, l(2)DTS513[1]* | Bloomington Drosophila Stock Center | BDSC: 5094 |
| *D. melanogaster*: *capu[EE] cn[1] bw[1]/CyO* | Bloomington Drosophila Stock Center | BDSC: 8788 |
| *D. melanogaster*: *P{pUAST-capu.mCherry}* | This study | N/A |
| *D. melanogaster*: *sn[3]* | Bloomington Drosophila Stock Center | BDSC: 113 |
| *D. melanogaster*: *w[\*]; P{w[+mW.hs]=FRT(w[hs])}G13 ena[210]/CyO* | Bloomington Drosophila Stock Center | BDSC: 25404 |
| *D. melanogaster*: *w[\*]; P{w[+mW.hs]=FRT(w[hs])}G13 tsr[N121]/CyO* | Bloomington Drosophila Stock Center | BDSC: 9109 |
| *D. melanogaster*: *y[1] w[\*]; P{w[+mC]=tubP-GAL80}LL10 P{ry[+t7.2]=neoFRT}40A/CyO* | Bloomington Drosophila Stock Center | BDSC: 5192 |
| *D. melanogaster*: *w[\*]; P{w[+mW.hs]=GawB}smid[C161]/TM6B, Tb[1]* | Bloomington Drosophila Stock Center | BDSC: 27893 |
| *D. melanogaster*: *y[1] w[\*]; Pin[Yt]/CyO; P{w[+mC]=UAS-mCD8::GFP.L}LL6* | Bloomington Drosophila Stock Center | BDSC: 5130 |
| *D. melanogaster*: *w[\*]; P{w[+mC]=UASp-GFP.Act5C}2-1* | Bloomington Drosophila Stock Center | BDSC: 9258 |
| *D. melanogaster*: *UAS-mCD8-Cherry/TM3* | Provided by Takashi Suzuki | N/A |
| *D. melanogaster*: *P{w[+m\*]=GAL4}5-40 P{w[+mC]=UAS-Venus.pm}1 P{w[+mC]=SOP-FLP}42; P{w[+mC]=tubP-GAL80}LL10 P{ry[+t7.2]=neoFRT}40A / CyO* | Kyoto Stock Centre | DGRC: 109947 |
| *D. melanogaster*: *P{w[+m\*]=GAL4}5-40 P{w[+mC]=UAS-Venus.pm}1 P{w[+mC]=SOP-FLP}42; P{w[+mW.hs]=FRT(w[hs])}G13 P{w[+mC]=tubP-GAL80}LL2 / CyO* | Kyoto Stock Centre | DGRC: 109948 |
| *D. melanogaster*: *w[\*]; P{w[+mW.hs]=FRT(w[hs])}G13* | Kyoto Stock Centre | DGRC: 106602 |
| *D. melanogaster*: *w[1118]; P{ry[+t7.2]=neoFRT}40A/CyO; P{ry[+t7.2]=neoFRT}80B* | Bloomington Drosophila Stock Center | BDSC: 8215 |
| Software and Algorithms |  |  |
| TREES toolbox | https://www.treestoolbox.org/ | N/A |
| MATLAB 2017b | https://se.mathworks.com/products/matlab | N/A |
| ImageJ | https://imagej.net/ | N/A |
| Prism7.0 (GraphPad) | https://www.graphpad.com/ | N/A |

**Table 2.** Genotypes.

| ABBREVIATION | GENOTYPE | FIGURE and PANNEL |
| --- | --- | --- |
|  | *P{w[+mC]=UASp-GFP,Act5C}2-1 / + ; P{w[+mW.hs]=GawB}smid[C161], UAS-mCD8-Cherry / +* | Figure 2, Movie S1 |
| control 40A | *P{w[+m\*]=GAL4}5-40 P{w[+mC]=UAS-Venus.pm}1 P{w[+mC]=SOP-FLP}42 ; P{w[+mC]=tubP-GAL80}LL10 P{ry[+t7.2]=neoFRT}40A ; P{w[+mC]=tubP-GAL80}LL10 P{ry[+t7.2]=neoFRT}40A / P{ry[+t7.2]=neoFRT}40A* | Figure 1A-E,H, 3B,C, 4B, 5D,E, 6D, S2A-C, S3D |
| arpc1 q25sd (FBal0008422) | *P{w[+m\*]=GAL4}5-40 P{w[+mC]=UAS-Venus.pm}1 P{w[+mC]=SOP-FLP}42 ; P{w[+mC]=tubP-GAL80}LL10 P{ry[+t7.2]=neoFRT}40A ; P{w[+mC]=tubP-GAL80}LL10 P{ry[+t7.2]=neoFRT}40A / Arpc1[Q25sd] P{ry[+t7.2]=neoFRT}40A* | Figure 3B,C, 4B, 5D,E, 6D, S2B,C, S3D |
| control G13 | *P{w[+m\*]=GAL4}5-40 P{w[+mC]=UAS-Venus.pm}1 P{w[+mC]=SOP-FLP}42 ; P{w[+mW.hs]=FRT(w[hs])}G13 P{w[+mC]=tubP-GAL80}LL2 / P{w[+mW.hs]=FRT(w[hs])}G13* | Figure 1A-E,H, 3D,E, 4B, 5B,C, 6E,G, S2A-C, S3E,G, S4 |
| ena 210 (FBal0031206) | *P{w[+m\*]=GAL4}5-40 P{w[+mC]=UAS-Venus.pm}1 P{w[+mC]=SOP-FLP}42 ; P{w[+mW.hs]=FRT(w[hs])}G13 P{w[+mC]=tubP-GAL80}LL2 / P{w[+mW.hs]=FRT(w[hs])}G13 ena[210]* | Figure 3D,E, 4B, 5B,C, 6E, S2B,C, S3E, |
| twinstar N121 (FBal0177372) | *P{w[+m\*]=GAL4}5-40 P{w[+mC]=UAS-Venus.pm}1 P{w[+mC]=SOP-FLP}42 ; P{w[+mW.hs]=FRT(w[hs])}G13 P{w[+mC]=tubP-GAL80}LL2 / P{w[+mW.hs]=FRT(w[hs])}G13 tsr[N121]* | Figure 3D,E, 4B, 5B,C, 6G, S2B,C, S3G, S4 |
| control | *P{w[+mW.hs]=GawB}smid[C161], P{w[+mC]=UAS-mCD8::GFP.L}LL6 / +* | Figure 1A-E,H, 3A,F-H, 4A,B, 5A,C, 6A-C,F, S1A-D, S2A-C, S3A-C,F |
| capu 1/EE (FBal0001537/ Fbal0045438) | *P{w[+mW.hs]=GawB}smid[C161], P{w[+mC]=UAS-mCD8::GFP.L}LL6 / + ; capu[EE] cn[1] bw[1] / capu[1] cn[1] bw[1]* | Figure 3F,G, 4B, 5A,C, 6B, S1C,D, S2B,C, S3B |
| spire 1/2F (FBal0016011/ Fbal0102386) | *P{w[+mW.hs]=GawB}smid[C161], P{w[+mC]=UAS-mCD8::GFP.L}LL6/+ ; spir[1] cn[1] bw[1] / b[1] pr[1] spir[2F] cn[1]* | Figure 3F,G, 4B, 5A,C, 6C, S1A,B, S2B,C, S3C |
| capu 1/+ | *P{w[+mW.hs]=GawB}smid[C161], P{w[+mC]=UAS-mCD8::GFP.L}LL6/+ ; capu[1] cn[1] bw[1] / +* | Figure 3H |
| spire 2F/+ | *P{w[+mW.hs]=GawB}smid[C161], P{w[+mC]=UAS-mCD8::GFP.L}LL6/+ ; b[1] pr[1] spir[2F] cn[1] / +* | Figure 3H |
| capu 1/spire 2F | *P{w[+mW.hs]=GawB}smid[C161], P{w[+mC]=UAS-mCD8::GFP.L}LL6/+ ; capu[1] cn[1] bw[1] / b[1] pr[1] spir[2F] cn[1]* | Figure 3H |
| singed 3 (FBal0015773) | *sn[3] / sn[3] ; P{w[+mW.hs]=GawB}smid[C161], P{w[+mC]=UAS-mCD8::GFP.L}LL6/+* | Figure 3F,G, 4B, 5A,C, 6G, S2B,C, S3F |
| UASspireHA | *P{w[+mW.hs]=GawB}smid[C161], P{w[+mC]=UAS-mCD8::GFP.L}LL6 / M{UAS-spir. ORF.3xHA}ZH-86Fb ; spir[1] cn[1] bw[1] / b[1] pr[1] spir[2F] cn[1]* | Figure S1A,B |
| UAScapu3MCherry | *P{w[+mW.hs]=GawB}smid[C161], P{w[+mC]=UAS-mCD8::GFP.L}LL6 / P{pUAST-capu.3M.mCherry} ; capu[EE] cn[1] bw[1] / capu[1] cn[1] bw[1]* | Figure S1C,D |

Single c3da clones (mosaic analysis with a repressible cell marker – MARCM) harbouring a null mutation in a component of the essential actin nucleator Arp2/3 complex component *arpc1* (**Figure 3B**), a strong hypomorphic allele of the actin polymerase *ena* (**Figure 3D**) or a loss of function allele of the actin severing factor *twinstar* (**Figure 3D**), as well as c3da neurons of larvae bearing a hypomorphic mutations for the actin bundler *singed* (**Figure 3F**), all showed reduced number of branches, as expected (**Figures 3C, E, G**; Gao et al., 1999; Nagel et al., 2012; Nithianandam and Chien, 2018; Stürner et al., 2019; Shimono et al., 2014).

In addition, mutants of the actin nucleators *spire* or *capu* displayed total numbers of branches in c3da neurons that were reduced by roughly a third (**Figure 3F, G**; **Figure S1)**. Thus, Spire and Capu represent novel regulators of c3da neuron morphology. The cooperation of Spire and Capu, is conserved across metazoa and extensively studied in *Drosophila* oocyte development (Dahlgaard et al., 2007). While individual *spire* or *capu* heterozygous mutants did not show any changes in morphology, their trans-heterozygous combination reduced the number of branches to a level comparable to that observed in the single homozygous mutants (**Figure 3H**). This suggests that Spire and Capu cooperate to define the number of c3da STBs.

Although each of these molecules has a distinct biochemical function in actin organisation all mutants showed a reduced number of branches (**Figure 3**) in c3da neurons. To reveal potential distinctions that might allow defining individual functions, we sought to define the morphology of wild-type c3da neurons and their STBs in greater detail.

### Distinctive roles of six actin-regulatory proteins on c3da dendrites

As a second step towards a quantitative description of c3da neuron dendrites and of the morphological effect of mutating individual AMPs, we identified a specific set of distinctive morphometric features for these neurons. We collected 28 general dendritic branching features (see **STAR**\***Methods**, **Table 3**) (Castro et al., 2020) and we used them to quantitatively describe c3da dendrites.

**Table 3.**
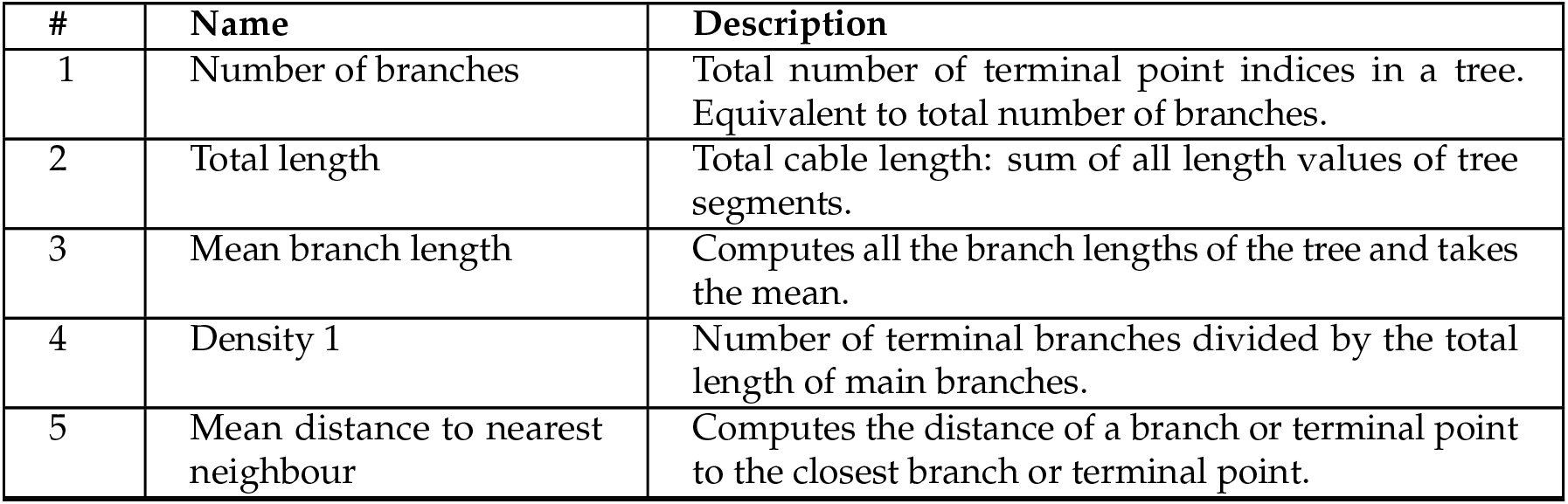
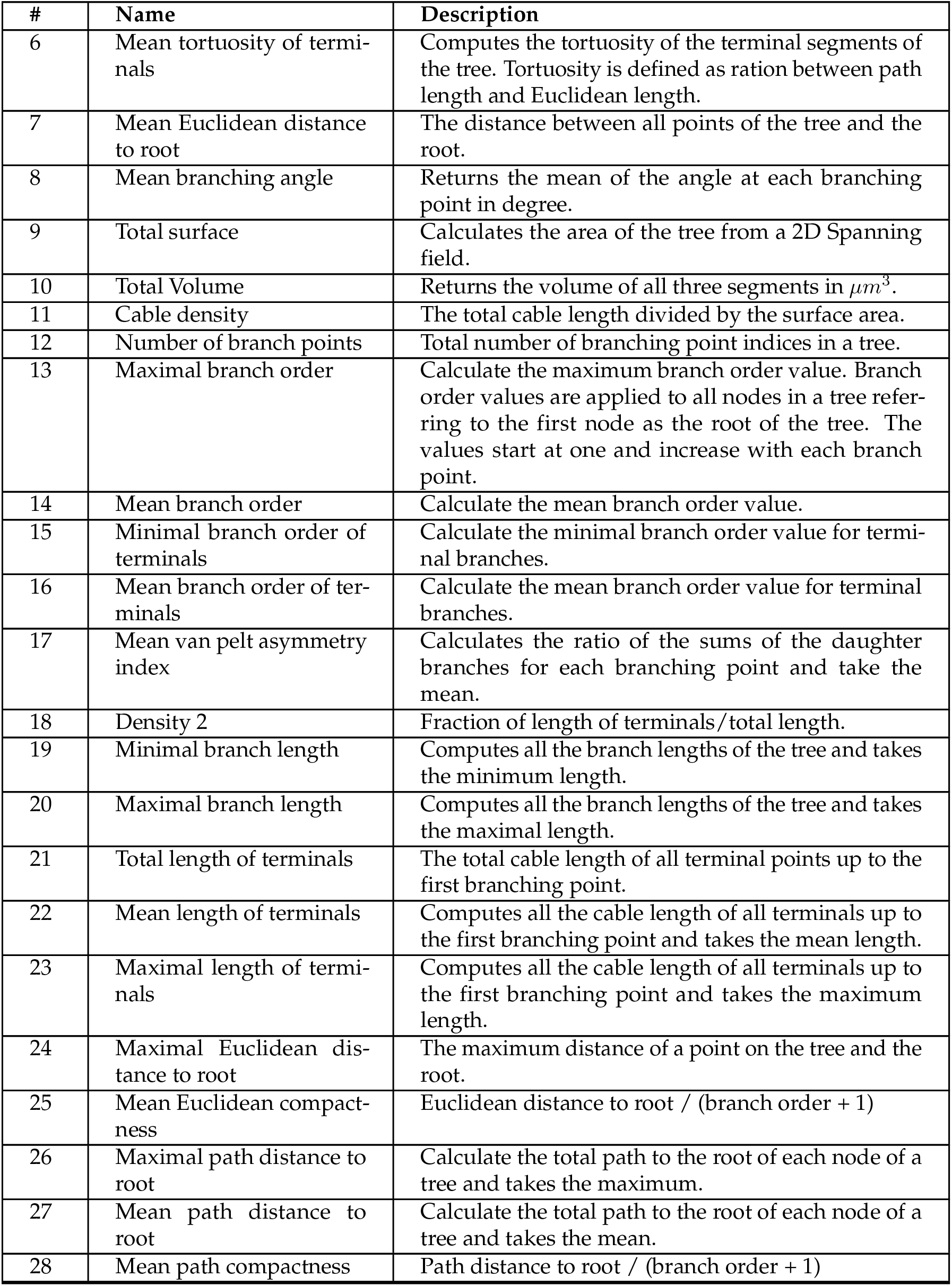
28 features with description.

| # | Name | Description |
| --- | --- | --- |
| 1 | Number of branches | Total number of terminal point indices in a tree. Equivalent to total number of branches. |
| 2 | Total length | Total cable length: sum of all length values of tree segments. |
| 3 | Mean branch length | Computes all the branch lengths of the tree and takes the mean. |
| 4 | Density 1 | Number of terminal branches divided by the total length of main branches. |
| 5 | Mean distance to nearest neighbour | Computes the distance of a branch or terminal point to the closest branch or terminal point. |
| 6 | Mean tortuosity of terminals | Computes the tortuosity of the terminal segments of the tree. Tortuosity is defined as ration between path length and Euclidean length. |
| 7 | Mean Euclidean distance to root | The distance between all points of the tree and the root. |
| 8 | Mean branching angle | Returns the mean of the angle at each branching point in degree. |
| 9 | Total surface | Calculates the area of the tree from a 2D Spanning field. |
| 10 | Total Volume | Returns the volume of all three segments in $\mu m^3$. |
| 11 | Cable density | The total cable length divided by the surface area. |
| 12 | Number of branch points | Total number of branching point indices in a tree. |
| 13 | Maximal branch order | Calculate the maximum branch order value. Branch order values are applied to all nodes in a tree referring to the first node as the root of the tree. The values start at one and increase with each branch point. |
| 14 | Mean branch order | Calculate the mean branch order value. |
| 15 | Minimal branch order of terminals | Calculate the minimal branch order value for terminal branches. |
| 16 | Mean branch order of terminals | Calculate the mean branch order value for terminal branches. |
| 17 | Mean van pelt asymmetry index | Calculates the ratio of the sums of the daughter branches for each branching point and take the mean. |
| 18 | Density 2 | Fraction of length of terminals/total length. |
| 19 | Minimal branch length | Computes all the branch lengths of the tree and takes the minimum length. |
| 20 | Maximal branch length | Computes all the branch lengths of the tree and takes the maximal length. |
| 21 | Total length of terminals | The total cable length of all terminal points up to the first branching point. |
| 22 | Mean length of terminals | Computes all the cable length of all terminals up to the first branching point and takes the mean length. |
| 23 | Maximal length of terminals | Computes all the cable length of all terminals up to the first branching point and takes the maximum length. |
| 24 | Maximal Euclidean distance to root | The maximum distance of a point on the tree and the root. |
| 25 | Mean Euclidean compactness | Euclidean distance to root / (branch order + 1) |
| 26 | Maximal path distance to root | Calculate the total path to the root of each node of a tree and takes the maximum. |
| 27 | Mean path distance to root | Calculate the total path to the root of each node of a tree and takes the mean. |
| 28 | Mean path compactness | Path distance to root / (branch order + 1) |

The combination of just seven of these 28 features accurately described the differences between the AMP mutant c3da morphologies (**Figure 4**). The total length of the dendrites together with the Euclidean distance of terminal points to the root represent the overall organisation of the tree; the mean length defines mean branch length distribution; the further parameters describe the distribution (density of terminals), the spreading (mean distance to nearest neighbouring terminal points and mean angle between branches) and the straightness of the terminal branches (tortuosity of branches) (**Figure 4A**). Here, the terminal branches are defined as all branches with a termination point, independently of their length.

**Fig 4.**
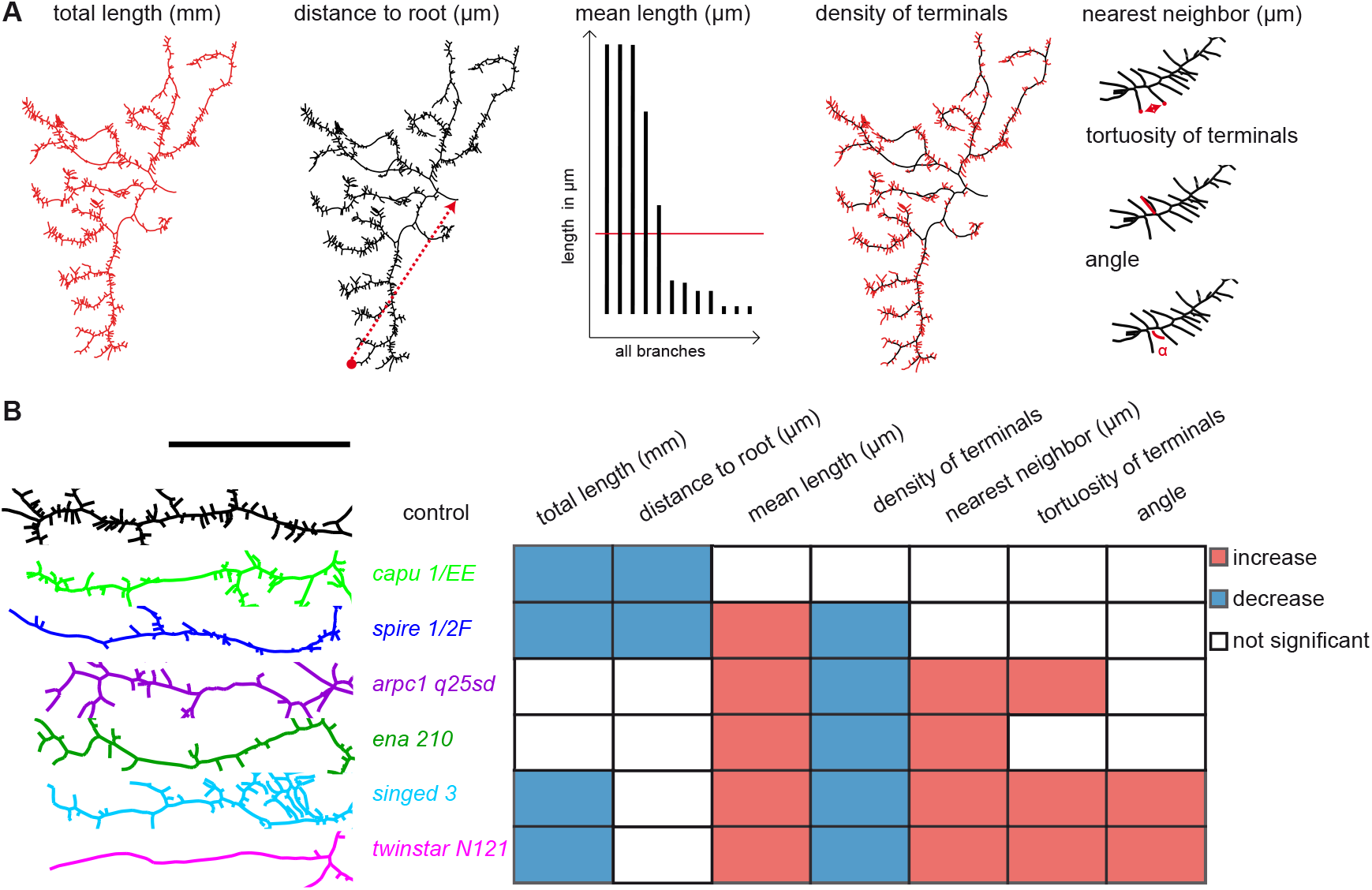
Features of dendritic tree structure in c3da neurons. **A**, Illustration of the seven morphometric measures defining c3da neuronal morphologies: total length of the dendritic tree in *mm* (total length), the mean Euclidean distance of terminal points to the soma in *μm* (distance to the root), the mean length of all branches in *μm* (mean length), the density of terminal branches along the length of main branches (black = main branches, red = terminal branches) (density of terminals), the distance of terminal points to the nearest neighbouring terminal point in *μm* (nearest neighbor), the mean tortuosity of branches (tortuosity of terminals), the mean angle between branches (angle). **B**, Image of one main branch with STBs of the control and each mutant in corresponding colours. Next to it, graphic representation of the seven morphometric measurements for each mutant versus corresponding controls. Blue for a significant decrease and red for a significant increase. *n* = 8 neurons from individual larvae per genotype. See **Figure S3** for complete graphs, **Table 2** for genotypes, **Table 3** for morphometric measures.

All mutants analysed displayed a reduction in the number of branches (**Figure 3**). In addition, *spire* and *capu* mutants showed a reduced total length and reduced distances to the root (**Figure 4B**). Indeed, *capu* and *spire* mutant trees were smaller and had most of their branches shifted closer to the cell soma (**Figure 4B**). While terminal branches properties seemed otherwise unaffected in *capu* mutants, the length of the main branches was reduced insuring a wild-type density of terminal branches along the main branches. Thus, Capu seems to promote branching and elongation of c3da dendrites. *Spire* mutants displayed instead an increase in mean length of branches and decreased density of terminal branches (**Figure 4**), suggesting that Spire, though involved in both, might promote branching over elongation (see **Figure 4B**).

The Arp2/3 complex is important for branch formation in all da neuron classes (Stürner et al., 2019). In the *arpc1* mutant c3da neurons this loss of branches was compensated by an increase in mean length to such an extent that the total length of the dendritic tree was not altered (**Figure 4B**). The reduced number of terminals and longer branches correlated with a decrease in the density of terminals. Moreover, the terminal branches of *arpc1* mutants were more spread out resulting in larger distances between neighbouring terminal points (**Figure 4B**). These data are consistent with a major role for Arp2/3 in the initiation of branching (Stürner et al., 2019).

*Ena* encodes a substrate of the tyrosine kinase *Abl* facilitating actin polymerisation (Damiano-Guercio et al., 2020; Brühmann et al., 2017). Ena plays a role in the elongation of lateral branches in dendrites of all classes of da neurons in the dorsal cluster (Gao et al., 1999). C4da neurons displayed dendrite over-elongation and reduced branching in *ena* mutants (Dimitrova et al., 2008). Likewise, in *ena* mutant c3da neurons the loss of terminal branches was compensated in part by increasing mean branch length and overall branch spreading, measured as distance to nearest neighbour (**Figure 6B**). Thus, similarly to *arpc1* mutants, *ena* mutant trees seem to counterbalance the loss of STBs by extending longer branches, pointing to a role of Ena in promoting branching over elongation (**Figure 4**).

**Fig 5.**
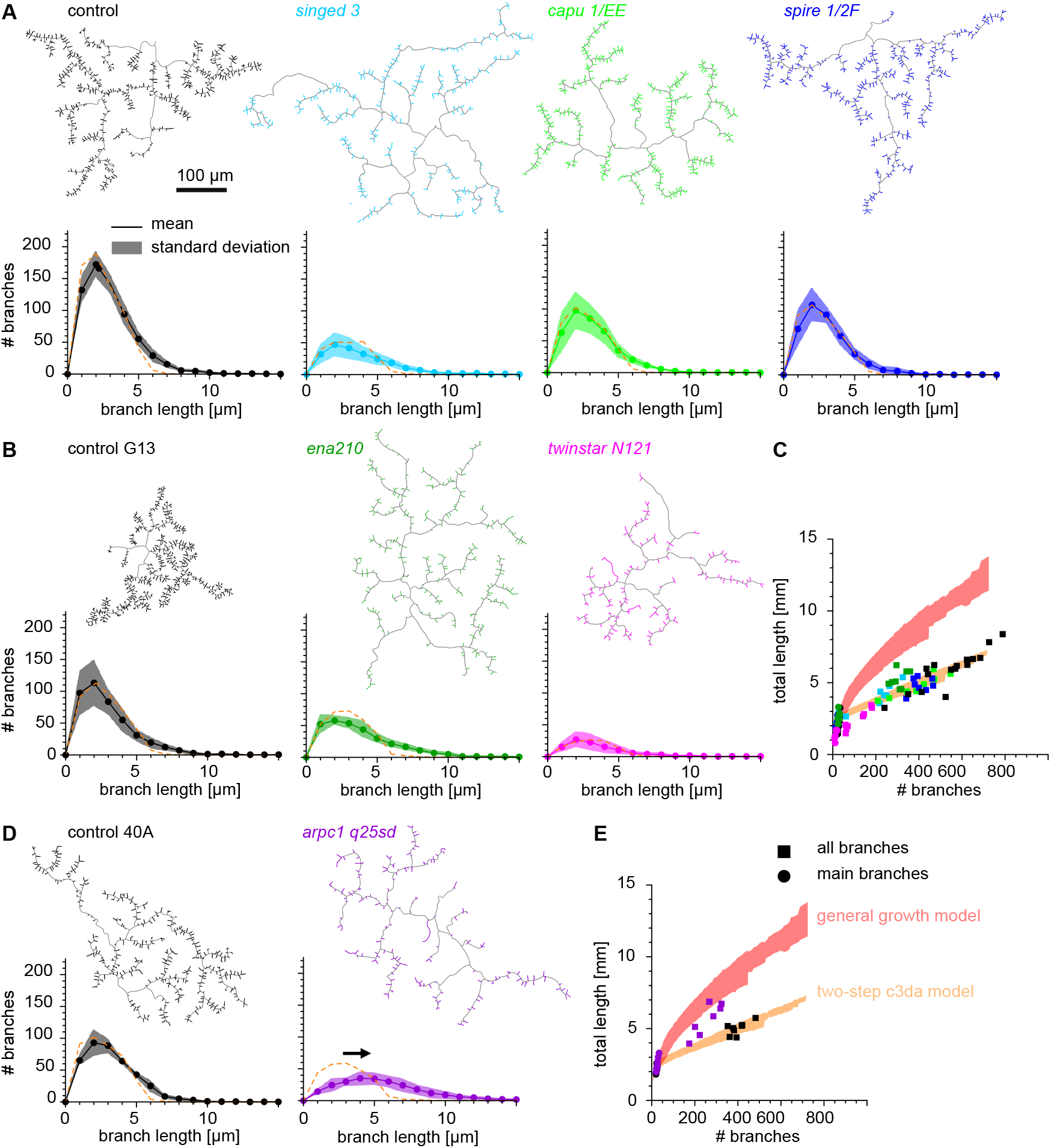
Modelling AMP mutant dendritic tree morphology using the c3da neuron model. **A, B, D**, The two-step c3da model applied to control and mutant dendritic trees; the STBs are represented in the colour corresponding to the genotype. The distribution of branch lengths for all STBs is shown underneath each neuron tracing. Distributions from the model in orange dashed line and distributions from real dendrites with respective colour corresponding to the genotype (see **Table 2** for genotypes). Arrow is pointing to the shift observed in the *arpc1* mutant. **C, E**, The real dendritic trees in coloured dots (only the main branches) and in coloured squares (all branches) are plotted with total length in *mm* to total number of branches. The trajectory for the c4da model is shaded red, the trajectory for the c3da model is shaded in orange.

In hypomorphic mutants for the actin bundling factor *singed* the total length reduction of c3da dendrites was not compensated by the increased mean length (**Figure 4B**) as it is instead the case in null *singed* mutants (Nagel et al., 2012). The branches were more spread out, with increased distance between neighbouring branches and decreased density of terminal branches. The few branches that were left had an increased tortuosity, as shown previously (Nagel et al., 2012), and were more spread with larger branching angles (**Figure 4B**). These data are consistent with Singed / fascin’s role in defining number and properties of the STBs (Nagel et al., 2012).

The dendrites of c3da neurons mutant for the severing protein *twinstar* display the most severe reduction in branch number (**Figure 3D,E**) that was not compensated by increased branch length yielding smaller trees (**Figure 4B**). The density of terminal branches was decreased and the branches left were more spread out, with increased distance between neighbouring branches and increased branching angle (**Figure 4B**). The naked main branches were more tortuous (**Figure 4B**). These data are consistent with a major role of Twinstar / cofilin in branch formation, although some terminal branchlets were still present in these mutants, typically close to the cell body.

Taken together, parallel evaluation of six AMP mutants pinpointed the seven morphometric features of c3da neurons that were necessary to describe differences in dendrite morphology between these AMP mutants, suggesting these might be key features of dendrite elaboration controlled by actin. Each of the AMP affected the organisation of the c3da neurons in characteristic ways hinting to specific roles during dendrite elaboration.

### The two-step c3da model can be applied to AMP mutant trees

Does the neuron still grow with the same core rules that we established for the wild-type c3da dendritic trees even in the AMP mutants and, if so, can we predict the morphology of mutant dendritic trees? To resolve this question, we used our two-step computational model to replicate the altered morphologies of the six AMP mutants.

We found that distributions of terminal branch lengths in *singed, spire, capu, ena* and *twinstar* mutants (**Figure 5A,B,C**) were modelled adequately with the two-step c3da model, given their respective dendrite field areas and the total number of branches obtained from the real data of each individual mutant tree (**Figure 1**). When comparing the distribution of terminal branch lengths obtained from the model (orange dashed line), they aligned with the distribution obtained from real dendritic trees (**Figure 5A,B**). Moreover, the scaling relations in real dendritic trees of the different mutants corresponded well to the c3da model trajectories in dark orange obtained previously in **Figure 1** (**Figure 5C**). Thus, the two-step c3da model replicated branching statistics for these mutants without requiring any modifications of the parameters established for the wild type, i.e. none of the core growth rules used to build the two-step c3da growth model were altered in these mutants.

However, the *arpc1* mutant dendritic trees could not be fully modelled with this two-step c3da model. While the spatial distribution of the main branches in the synthetic trees revealed by Sholl analysis resembled the wild type (**Figure S2A,B**) the distribution of terminal branch length in the model predicted shorter branches than observed in the real *arpc1* mutant dendritic trees (**Figure 5D**). Thus, the two-step c3da model did not replicate the *arpc1* mutant trees in their distribution of lengths of STBs nor in the correlation of total length to branch number (**Figure 5E**). The resulting scaling relationships as well as the longer terminal branch lengths indicated that *arpc1* mutant trees might lie somewhere between the c4da and the new suggested c3da wild type model (**Figure 5D,E**).

Taken together, based on the spanning area of a dendrite, its total length and the distribution of branches, our new two-step c3da model was able to predict aspects of the dendritic tree morphology of five out of six AMP mutants that we investigated. The c3da model does not include a detailed description of the morphological properties of STBs. Nonetheless, the wild-type c3da model directly predicted the length distributions of the STBs of five AMP mutants. This indicates that these five AMPs do not affect the core rules that define c3da dendrite distribution. In case of the *arpc1* mutant dendritic tree, however, the dendrite defect cannot be accurately modeled, suggesting that a core aspect of dendrite organisation is altered in this mutant.

### Contribution of individual actin-modulatory proteins to complex branchlet dynamics

There are different ways in which the reduction of dendritic branches and the specific alterations observed in the mutants could arise. For instance, reduction of branches could be caused by defects in dendrite maintenance, increased dendrite retraction or by reduced branch formation. To gain a clearer understanding of the origin of the morphological alterations observed in the different AMP mutants we performed time-lapse analysis in live animals (see **STAR**\***Methods**). Immobilised larvae of late *2^nd^* instar stage were imaged every minute over 30*min.* To simplify the analysis, we down-sampled to trace only every fifth minute and tracked the STBs over time using a dedicated user interface (Baltruschat et al., 2020) and *ad hoc* scripts (`ui_tlbp_tree`) in the *TREES toolbox* (www.treestoolbox.org), enabling to compare the dynamics between animals and groups.

STBs were categorised into one of the following five groups: stable, new, extending, retracting and disappearing branches, depending on the dynamics observed between one time point and the following (**Figure 6A**). We additionally tracked the terminal and branch points to measure the velocity of extension and retraction of branches quantified as the travelled distance of the branch tip over time 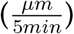.

The loss of *capu, spire* or *arpc1* led to a reduced number of newly forming branches (**Figure 6B– D**), suggesting that these actin nucleation factors are important for the very first step of branch formation, as previously already demonstrated for *arpc1* (Stürner et al., 2019). In addition, mutants of *spire* showed an increase in stable branches that was linked to a decrease in the number of extending, retracting and disappearing branches (**Figure 6C**). Thus, Spire displayed an additional role in branch dynamics, possibly linked to a function independent of Capu. The higher resolution of the time-lapse analysis in c3da neurons also suggested an additional, previously unrevealed, role for *arpc1* in promoting retraction and disappearance of branches, as both were decreased in the mutant condition (**Figure 6D**, Stürner et al., 2019).

Time-lapse imaging revealed an increase in branch extension and new formation of STBs in absence of *ena*, which suggests that Ena hinders formation or extension of STBs (**Figure 6E**). Consistently, there was a decrease in disappearing branches, indicating again that Ena could be limiting the characteristic dynamics of the STBs thereby promoting them to develop into long main branches (**Figure 6E**).

Singed / fascin supports the formation of unipolar actin filament bundles and is suggested to give filopodia the stiffness necessary for membrane protrusion (Vignjevic et al., 2006). Our improved time-lapse analysis revealed that this stiffness although required for the characteristic straightness of the STBs, does not facilitate the dynamical movement of the branchlet (**Figure 6F**). A reduction in the amount of Singed / fascin in the c3da neurons in fact led to an overall increase in dynamics, suggesting that tight unipolar bundling of actin is restricting the dynamics of the branchlet to provide this stiffness (**Figure 6F**).

**Fig 6.**
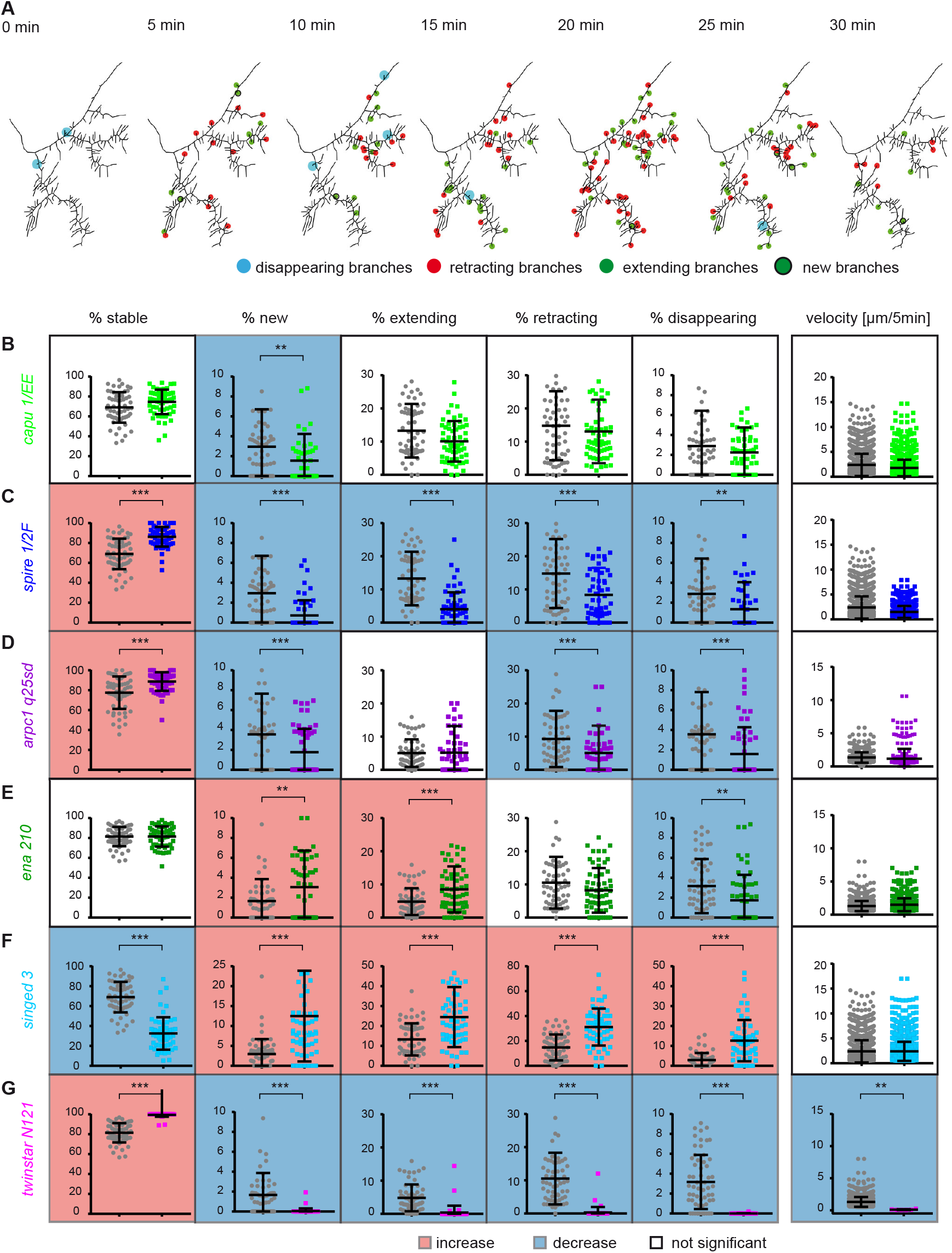
Time-lapse analysis of terminal dendritic branches of c3da neurons. **A**, Representative example of a tracing of a terminal region of a control c3da neuron over 30min in 7 steps of 5*min*. Terminal branches that disappeared (blue), retracted (red) extended (green), or newly formed (green with black ring) from one time point to the next are marked with a dot in the corresponding colour. **B–G**, Percentage of terminal branches that were stable, new, extending, retracting or disappearing between timepoints over 30min of time-lapse for each mutant versus corresponding control (grey/black). Average velocity of a terminal branch, quantified as the average change in length (extension + retraction) in 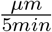 (corrected p values * is *p* < 0.05, ** is *p* < 0.01 and *** is *p* < 0.001). Tł graph background is highlighted in blue for a significant decrease and in red for a significant increase *n* = 10 neurons from individual larva per genotype (see **Table 2** for genotypes).

In partial agreement with recent data (Nithianandam and Chien, 2018) obtained by RNAi, the loss of *twinstar* showed almost no newly forming, extending, retracting or disappearing branches in distal regions of the dendritic tree (**Figure 6G**). Thus, STB formation is very limited without actin remodelling through *twinstar* and branch dynamics is strongly reduced. However, areas of the dendritic tree closer to the cell body still displayed some STBs, which had normal dynamics properties (see **Figure S4**). This might be explained with residual Twinstar / cofilin protein in the mutant neuron, sufficient to guarantee branchlet formation and dynamics in these segments.

Taken together, by examining the loss of individual AMPs in the same dendritic branchlet in a comparative way together with a detailed quantitative description of dynamics alterations in the AMP mutants we could make a first attempt at understanding how together these AMPs define the specific dynamics of c3da STBs.

## Discussion

Neurons develop their dendrites in tight relation to their connection and computation requirements (Poirazi and Papoutsi, 2020). Thus, dendrite morphologies display sophisticated type-specific patterns. From the cell biological and developmental perspective this raises the intriguing question at which level different neuronal types might use shared mechanisms to assemble their dendrites. And conversely, how are specialised structures achieved in different neuronal types? To start addressing this core question of neuronal cell biology, we tightly combined a computational and a cell biological approach. We found by modelling the morphology of the *Drosophila* larva c3da neurons that two distinct growth programs, are required to achieve models that faithfully reproduce the dendrite organisation of those neurons. The model singles out the STBs of c3da neurons that are also molecularly identifiable as specific structures. By combining time-lapse *in vivo* imaging and genetic analyses, we shed light on the machinery that controls the dynamic formation of those branchlets.

### A molecular model of branchlet dynamics

The complex interplay of AMPs generates highly adaptive actin networks. In fact, in contrast to earlier unifying models, it is now clear that even the same cell can make more than one type of filopodium-like structure (Bilancia et al., 2014; Barzik et al., 2014). Here we characterised the effect of loss of six AMPs on the morphology and dynamics of one specific type of dendritic branchlet, the STBs of c3da neurons. With this information, we delineate a molecular model for branchlet dynamics *in vivo* in the developing animal (**Figure 7**). Similar approaches to model the molecular regulation of actin in dendrite filopodia have been taken recently for cultured neurons (Marchenko et al., 2017). In comparison to those, we rely directly on the effect of loss of individual AMPs *in vivo*. The advantage of the present *in vivo* approach is that it preserves the morphology, dynamics and adhesive properties of the branchlets and non cell-autonomous signals remain present.

**Fig 7.**
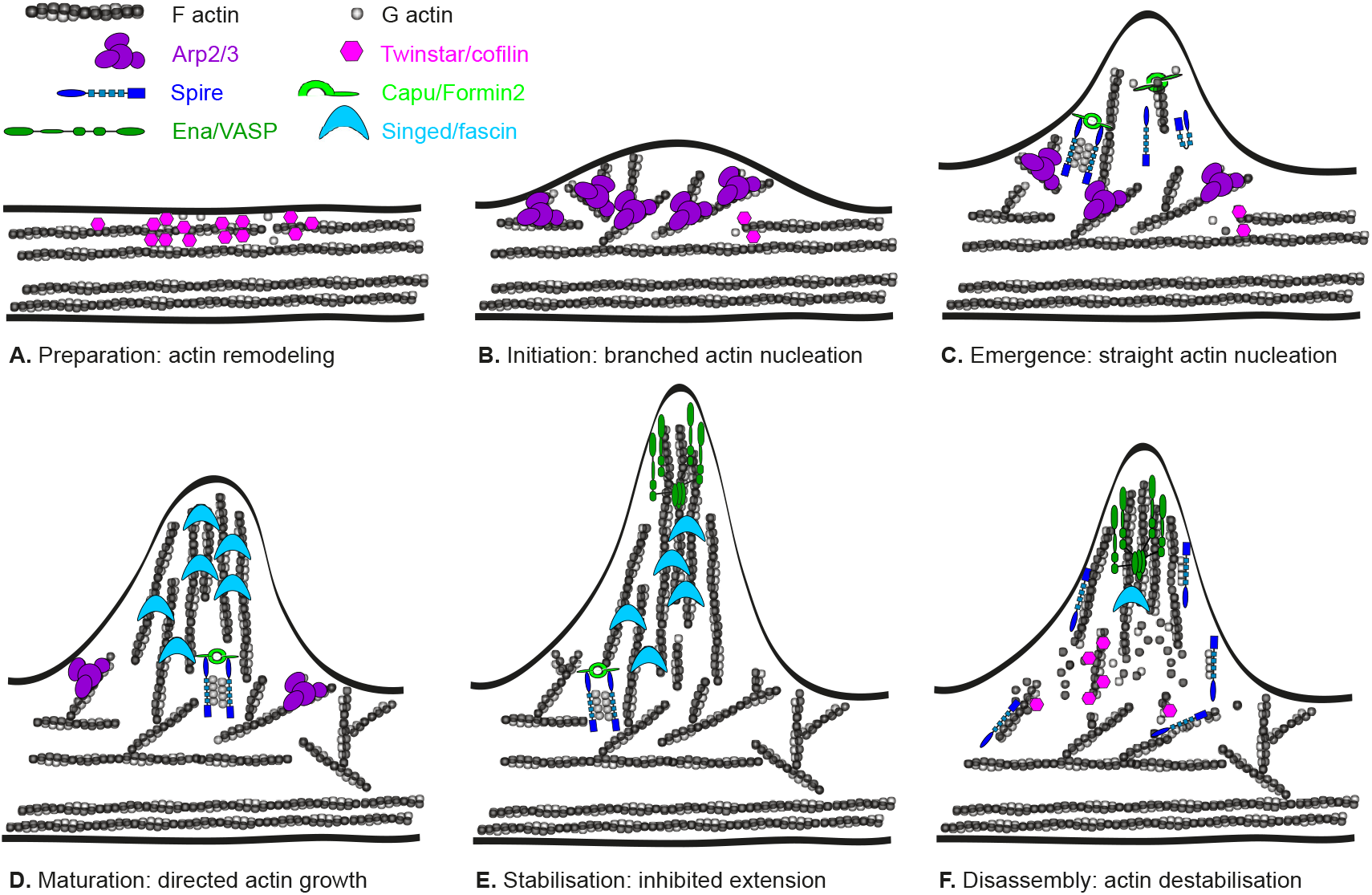
A theoretical model of dendritic branch dynamics. **A**, Actin remodelling and availability of a pool of monomeric actin (G actin), provided by Twinstar / cofilin, is a prerequisite for the formation of new filamentous actin structures (F actin). **B**, Membrane protrusion requires a branched actin network at the base, mediated by the actin nucleation complex Arp2/3. **C**, Straight actin filaments, nucleated by Spire and Capu / Formin2 together, push out the membrane before **D**, the actin filaments can be bundled by Singed / fascin, to restrict their dynamics and give them their characteristic angle and shape. **E**, The presence of Singed / fascin facilitates the binding of Ena / VASP, which limits the mature terminal branchlet from extending further. **F**, Terminal branches regularly retract and can disappear completely, facilitated by Ena / VASP and Spire that can destabilise the filaments.

The combination of our FRAP experiments pointing to fluorescence recovery at the distal tip of an extending branchlet and the localisation of Singed / fascin on the extending terminal branchlets (Nagel et al., 2012) strongly suggested that actin is organised in a tight bundle of mostly uniparallel fibres with the barbed end oriented distally in c3da neurons. This organisation is thus very different from what that of dendritic filopodia of hippocampal neurons in culture (Portera-Cailliau et al., 2003; Svitkina et al., 2010; Marchenko et al., 2017). The actin filaments in the bundle appear to be particularly stable in the c3da neuron STBs as the actin turnover that we revealed by FRAP analysis was 4 times slower than what would be expected in dendrite spines of hippocampal neurons *in vitro* (Star et al., 2002; Zito et al., 2004) and 20 fold slower than in a lamellipodium of melanoma cells *in vitro* (Lai et al., 2008). It is nonetheless in line with previous data on stable c3da neuron terminal branchlets (Andersen et al., 2005) and with bundled actin filaments of stress fibers of human osteosarcoma cells (Hotulainen and Lappalainen, 2006). We observed treadmilling, similarly to that of filopodia at the leading edge (Mallavarapu and Mitchison, 1999). The retrograde flow rate is 30 times slower than what has been reported for filopodia in hippocampal cells (Chazeau et al., 2015) and comparable to rates observed for developing neurons in culture lacking the mammalian homologues of Twinstar, ADF / cofilin (Flynn et al., 2012). Slower actin kinetics within the STBs could therefore mean that Twinstar / cofilin is not present within the STBs and only essential for a preliminary step of STB formation. Alternatively, the slow kinetics might be related to the fact that we are imaging neurons differentiating in the complex 3D context of a developing animal. Recent quantification of actin treadmilling in a growth cone of hippocampal neurons in 3D culture, though, did not produce differences with classical 2D culture models (Santos et al., 2020).

In summary, in c3da STBs actin is organised in uniparallel bundles with their barbed ends pointing distally and the filaments display characteristic slow dynamics of polymerisation and of treadmilling.

What is the role of AMPs in defining the organisation of the actin filaments in the c3da STBs? The alterations of dendrite and STB morphology and dynamics caused by loss of individual AMPs function reported here can be combined with preceding molecular knowledge about these conserved factors to produce a hypothetical model of the actin regulation underlying STBs dynamics (**Figure 7**). Dendrite structure and time-lapse imaging point to an essential role of Twinstar / cofilin for the initiation of a branchlet, in agreement with previous literature (Nithianandam and Chien, 2018) (**Figure 7**). *Drosophila* Twinstar / cofilin is a member of the ADF / cofilin protein family, with the functional capacity of severing actin filaments, but with poor actin filament depolymerising activity (Shukla et al., 2018; Gunsalus et al., 1995). We thus propose that Twinstar / cofilin can induce a local fragmentation of actin filaments that can then be used as substrate by the Arp2/3 complex. In fact, in c4da neurons, Arp2/3 localises transiently at the site where the branchlets will be formed and its presence strongly correlates with the initiation of branchlet formation (Stürner et al., 2019). Previous and present time-lapse data pointed to the role of Arp2/3 in the early phases of branchlet formation (Stürner et al., 2019) (**Figure 6D**). Thus, we suggest that localised activity of Arp2/3 generates a first localised membrane protrusion (Mogilner and Oster, 1996).

Since the localisation of Arp2/3 is transitory (Stürner et al., 2019), we have interrogated the role of additional potential actin nucleators in this context. The formin family proteins regulate both the microtubule and the actin cytoskeleton in neurons (Szikora et al., 2017). Formins are associated with a variety of neurological disorders, though causative evidence remains elusive (Boyer et al., 2011; Lybaek et al., 2009; Ercan-Sencicek et al., 2015; Schymick et al., 2007). From an RNAi-supported investigation of the role of formins for da neuron dendrite morphology, we identified Capu as a potential modifier of c3da STBs (Stürner et al., 2019). Capu displays complex interactions with the actin nucleator Spire during oogenesis, involving cooperative and independent functions of these two molecules (Dahlgaard et al., 2007). An increase in Spire levels correlates with a smaller dendritic tree and inappropriate, F-actin-rich and shorter dendrites in c4da neurons (Ferreira et al., 2014). In our hands, though, loss of Spire function did not yield a detectable phenotype in c4da neurons. In c3da neurons, we found that Capu and Spire support the formation of new branchlets and display a strong genetic interaction in the control of branchlet number (**Figure 3H**). We thus suggest that they cooperatively take over the nucleation of linear actin filaments possibly producing the bundle of uniparallel actin filaments. However, the range of effects of loss of Spire function is broader than that of Capu, suggesting additional independent functions of Spire. Spire itself is a weak actin nucleator (Quinlan et al., 2007). We thus surmise that its Capu-independent properties are not related to nucleation. In the context of c3da STBs, Spire seems to promote branch dynamics. While we do not have a clear indication for the molecular mechanisms supporting this function, an actin severing activity of Spire was reported *in vitro* (Bosch et al., 2007). Although *in vivo* evidence for this function is lacking, the role of Spire on STB dynamics appears to be consistent with favouring actin destabilisation or actin dynamics (**Figure 6C**).

Singed / fascin bundles actin filaments specifically in the c3da neuron STBs and gives these branches their straight conformation (Nagel et al., 2012). The localisation of Singed / fascin in the c3da STBs correlates with their elongation (Nagel et al., 2012) and our present data point to Singed / fascin as a key regulator of STB dynamics and morphology. While the complete loss of *singed* function suppressed dynamics (Nagel et al., 2012), the mild reduction in protein level analyzed here led to more frequent branchlets elongation and retraction. Further, the branchlets extended at wrong angles and displayed a tortuous path. Singed / fascin controls the interaction of actin filament bundles with Twinstar / cofilin and can enhance Ena-mediated binding to barbed ends (Bachmann et al., 1999; Winkelman et al., 2014). Thus, in addition to generating mechanically rigid bundles (Mogilner and Rubinstein, 2005), it can modulate actin dynamics by regulating the interaction of multiple AMPs with actin. We speculate that the retraction and disappearance of the STB could be due to Singed / fascin dissociating from the actin filaments possibly in combination with Spire or Twinstar / cofilin additionally severing actin filaments (**Figure 7)**.

Ena is important for restricting STB length and it inhibits new formation and extension of STBs. This appears to be a surprising function for Ena, which is in contrast to its expected role in promoting actin filament elongation by antagonising actin filament capping and by processive actin elongation (Barzik et al., 2005; Bear and Gertler, 2009; Krause et al., 2002; Breitsprecher et al., 2011; Hansen and Mullins, 2010; Pasic et al., 2008), or to its capacity of supporting the activation of the WAVE regulatory complex (Chen et al., 2014). Similarly to what we previously reported for *ena* mutant c4da neurons, we observe a balance between elongation and branching also in c3da neurons (Dimitrova et al., 2008). In *Drosophila* macrophages, Ena was shown to associate with Singed / fascin within lamellipodia (Davidson et al., 2019). Along the line of these recent data, we suggest that Ena might have a similar function in the formation of the STBs and could closely cooperate with Singed to form tight actin bundles that slow down STB elongation.

### Quantitative analysis of neuronal morphology

The investigation of morphological parameters in combination with genetic analysis has proven extremely powerful to reveal initial molecular mechanisms of dendrite differentiation (Gao and Bogert, 2003). Early studies, though, have been limited in the description power of their analysis concentrating on just one or two parameters (e.g. number of termini and total dendrite length). This limitation has been recognised and addressed in more recent studies (Nanda et al., 2018b; Kanaoka et al., 2019; Das et al., 2017; Wang et al., 2019; Sheng et al., 2018; Li et al., 2017).

A major outcome of our present work is the establishment of a powerful tool to compare quantitatively different mutant groups. A detailed tracing of neuronal dendrites of the entire dendritic tree or a certain area of the tree in a time-series with a subsequent automatic analysis allows a precise description of the mutant phenotypes. We additionally generated novel tools for extracting quantitative parameters of the dynamic behaviour of dendrite branches from time-lapse movies based on a novel branch registration software (Baltruschat et al., 2020). This time-lapse tool operates similarly as in (Sheng et al., 2018) and was developed in parallel to (Castro et al., 2020), with the advantage of having an automated quantification after registration which detects branch types and their dynamics. Moreover, the tool operates in the same framework as the tracing and morphological analysis. We make these tools available within the *TREES toolbox* (www.treestoolbox.org, Cuntz et al., 2010) and encourage their use to support comparative analysis among data sets.

Our present data support a molecular model of dendrite branchlet formation and dynamics. They demonstrate that computational analysis can support a detailed quantification, revealing differences among even similar mutant phenotypes. Importantly, it can help to trace back the function of a protein and elicit new insights into complex molecular phenomena.

### Specialised growth programs to refine individual neuron type dendrite morphology

What are the fundamental principles that define dendrite elaboration and which constraints need to be respected by neurons in establishing their complex arbours? High-resolution time-lapse imaging together with digital reconstructions has pushed the quality of dendrite structure analysis, as discussed above. Here, we combined these tools with mathematical modelling to infer the growth rules underlying the establishment of a specific dendritic tree.

Models based on local or on global rules have been applied to reproduce the overall organisation of dendritic trees, including da neurons (Nanda et al., 2018b; Baltruschat et al., 2020; Castro et al., 2020). We based our c3da model on the fundamental organising principle that dendrites are built through minimising cable length and signal conduction times (Cuntz et al., 2007; Wen and Chklovskii, 2008; Cuntz et al., 2010; Baltruschat et al., 2020). This general rule for optimal wiring predicts tight scaling relationships between fundamental branching statistics, such as the number of branches, the total length and the dendrite’s spanning area (Cuntz et al., 2012). However, we observed that the characteristic STBs of c3da dendrites do not follow this scaling behaviour. Instead, we have shown that a second growth program must be postulated to account for their specific morphology. This is an interesting deviation from the general developmental growth model presented in Baltruschat et al. (2020).

Here, we found that c3da neurons respect the general growth model when stripped of all their STBs. This points to a basic layer of organisation that is shared among different types of neurons. A second, specialised step had to be applied to add the STBs to this basics structure, respecting their number, total length and distribution. Interestingly, the regularity index *R*, a recent branching statistic that is based on the nearest neighbour distances of terminal points in dendrites, had singled out c3da neurons for their comparably small *R* values, indicating a high clustering of branches presumably due to the c3da characteristic STBs (Anton-Sanchez et al., 2018). The two-step model used in this work suggests that while main dendritic trees have common growth rules that are balancing between efficiency and precision, the dendritic specialisations of any neuronal cell type need to be studied carefully, since the details do not necessarily have the same constraints. This view is compatible with findings in a companion paper where functional constraints shape the dendrites of c1da neurons in a specialised branch retraction phase additionally to the general growth phase that guarantees optimal wiring (Castro et al., 2020).

In our c3da dendrite model the resulting synthetic morphologies resemble the real dendritic trees including those of 5 out of the 6 ARP mutant dendritic trees without any changes to the model parameters. In addition to providing a new insight into how specialised dendritic trees are built, the model enables quantitative predictions for future questions.

In conclusion, we hypothesise that neuronal dendrites are built based on common, shared growth programs. An additional refinement step is then added to this scaffold, allowing each neuron type to specialise based on its distinctive needs in terms of number and distribution of inputs. In the exemplary case of the c3da neurons, we investigated molecular properties of these more-specialised growth programs and propose a first comprehensive model of actin regulation that explains the morphology and dynamics of branchlets.

## Acknowledgments

We are grateful to Dr. F. Bradke, Dr. G. Marchetti, Dr. K. Rottner, Dr. P. Soba, B. Schaffran and Dr. A. Ziegler for comments on the manuscript. This work was supported by a DFG grant (Teilprojekt, SPP 1464: Principles and evolution of actin-nucleator complexes) to G.T., a BMBF grant (No. 01GQ1406 — Bernstein Award 2013) to HC and a DFG grant (CU 217/2-1) to H.C.. We thankfully acknowledge Dr. E. Kerkhoff and A. Samol-Wolf for the Capu constuct. We would like to thank Dr. K. Rottner for discussion on the FRAP analysis. We thank R. Kerpen for great technical assistance throughout this work. The authors declare to have no competing financial interests.

## Author contributions

T.S., A.F.C., M.P., H.C., and G.T. designed the study. T.S. performed the experiments. T.S. and A.F.C. designed and analysed the time-lapse analysis. H.C. designed the growth models and performed the simulations. T.S., A.F.C., H.C., and G.T. wrote the paper.

## STAR*Methods

### METHODS DETAILS

#### Fly strains

Flies were reared on standard food in a 12*hr* light-dark cycle at 25°*C* and 60% humidity unless otherwise indicated.

A pUAST (Brand and Perrimon, 1993) containing a full-length Capu construct with a mCherry fluorescent tag (Q24120, 1059 aa) (kindly provided by Annette Samol-Wolf and Prof. Dr. Eugen Kerkhoff) was injected by BestGene Inc. (Chino Hills, CA, USA) to the 3^*rd*^ Chromosome.

#### Microscopy / Live imaging

For all of the imaging in this work living larvae were covered in Halocarbon oil, to allow oxygen exchange and immobilised between a cover slip and a glass slide. After imaging larvae were checked for vitality and set back on fly food, images taken from larvae that did not survive until hatching were excluded from the analysis. The larva were placed on their side to allow the imaging of the same lateral c3da neuron (ldaB) of the abdominal segment A5.

The FRAP experiments the same anterior portion of the ldaB neuron of late second instar larva were imaged with a LSM 800 Airyscan Microscope and a 63 × /1.40 oil objective **Figure 1A**. A 488*nm* for *GFP* and 561*nm* for *mCherry* line of an argon laser was used. The frame, including the ROI (tip of a branchlet), was imaged at least three times before bleaching. The laser was set to 90% maximal power for bleaching and 2% maximal power for imaging. Photo-bleaching was achieved with 10 iterations (scan speed at 3) of the region of interest. Imaging of the area was resumed immediately after photo-bleaching and continued every 30*sec* for at least ~ 300*sec*.

For **Figure 2** and **Figure 5** the entire dendritic tree of early third instar *Drosophila melanogaster* larvae were imaged with a LSM 780 Zeiss 40 × oil objective, the software used was ZEN 2010. One neuron was imaged per animal, 8 animals per genotype.

For the time-lapse series in **Figure 6** over 30*min* every 30*sec* was taken of an anterior portion of the ldaB neuron of late second instar larva with an Yokogawa Spinning-Disc on a Nikon stand (Andor, Oxford UK) with two back-illuminated EM-CCD cameras (Andor iXON DU-897) and a 60 × oil objective. One neuron was imaged per animal, 10 animals per genotype.

#### FRAP analysis

For the FRAP analyses w*; *theP{GawB}smidC 161/TM6B,Tb1* (B#27893) (Shepherd and Smith, 1996) was recombined with *UAS – mCD8 – Cherry/TM*3 (kindly provided by Takashi Suzuki) and crossed to w*; *P{UASp – GFP.Act5C*}2 – 1 (B# 9258).

A line analysis was conducted in the *ImageJ* software (version 1.52a) over time and space with a short macro that measures the intensity (*I_GFP_*, *I_mCherry_*) of each pixel of the two channels along the line over time. Moreover it tracks the extension of the branch along the line by comparing the intensity to an adjustable threshold (see script: `Analysis_FRAP_macro`). Background fluorescence intensities (*I_GF Pbg_*, *I_mCherrybg_*) taken from a region outside the cell were subtracted from each individual region and frame. The values were normalised to the average of 3 pre-bleach values (*I_N_*). Acquisition photo bleaching was determined by comparing the normalised *mCherry* signal (*I_mCherry_*) in the bleached area over time, the area seems unaffected by experimental bleaching as there is even an increase in *mCherry* signal over time. In **Figure 1D** the normalised GFP fluorescence 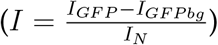 is visualised over time. Time point 0 (*t*_0_) was defined at the first time point after photo bleaching (after 2*min*) and the last time point as the *t*_∞_. The average halftime recovery was calculated 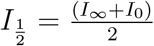 and the time point closest was defined as 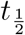. The average retrograde movement of actin (*M*) was quantified by drawing a line at the distance the pixel below a 30% Intensity threshold had from the originally bleached area toward the main branch. There is a very slow retrograde movement of 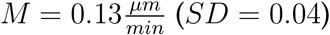.

#### Dendritic arbour analysis

Eight image stacks per genotype were manually reconstructed in 3D using the user interface `cgui_tree` of the *TREES toolbox* (www.treestoolbox.org) (Cuntz et al., 2010), an open source software package for *MATLAB* (Matworks, Natick, MA). A large palette of 28 branching statistics (**Table 3**) specifically for the c3da neurons were collected for each set of dendrite reconstructions using TREES toolbox functions. These branching statistics are aggregated in our new `features_c3_tree` function.

#### Time-lapse analysis

Ten image series per genotype were analysed. The single images of the 30*min* time series were manually reconstructed in 2D using the user interface `cgui_tree` of the *TREES toolbox* (www.treestoolbox.org) (Cuntz et al., 2010) every 5*min*. Then they were registered using the `ui_tlbp_tree` script as described in (Baltruschat et al., 2020) tracking terminal and branch points. The `eval_timelapse` script categorises the terminal branches into 5 groups: new branches that appear throughout the 30*min* and disappearing branches, branches with are extending or retracting and branches that do not change in length within a certain threshold. These numbers were divided by the total number of branches within the image frame. This allowed us to compare the different mutants and the branch dynamics independently of their difference in total branch number at the beginning of the imaging session. Moreover the `eval_timelapse` script computes the velocity of branch movement, as the average distance covered by a terminal branch over time (see `script_timelapse_analysis`). This analysis was developed in parallel to the time-lapse analysis in (Castro et al., 2020).

#### Statistical analysis

Data were analysed using *Prism 7.0* (GraphPad). Groups were compared using the Kruskal-Wallis test followed by Dunn’s post hoc test accordingly. Single comparisons between two groups were analysed using the two tailed Wilcoxon Signed Rank Test. For multiple comparisons with several features for each group the p values were controlled for false discovery rate by the adaptive method of Benjamini, Krieger and Yekutieli with a *Q*% of 3 (Benjamini et al., 2006) and controlled for statistical significance with the Holm-Sidak method (alpha of 0.05). Normal distribution of the dataset was confirmed using the Shapiro-Wilk and Kolmogorow-Smirnow normality test. The *p* values shown are all adjusted *p* values. (* is *p* < 0.05, ** is *p* < 0.01 and *** is *p* < 0.001).

#### Computational Modelling

The c4da neuron model was described previously in Baltruschat et al. (2020) and is provided there as a *TREES Toolbox* function `growth_tree`. Briefly, at each growth iteration, a new target is selected within the dendritic spanning area but far away from the existing tree. A parameter *k* determines the stochasticiy of the selection of the new target with a value of 0 referring to the target being as far as possible from the existing tree without any noise and 1 the target being chosen completely at random. A balancing factor *bf* weighs total cable length cost against mean path length to the soma (Cuntz et al., 2007, 2010). A parameter *radius* determines the outreach threshold that a new branch can grow to, restricting the area in which a target can be selected. This model was obtained from developmental growth iterations in time-lapse images and reproduces both the c4da morphology accurately as well as –though with different parameters-the morphology of a large number of dendrites from other cell types. The c4da model parameters were *k* = 0.45, *bf* = 0.225 and *radius* = 120*μm*. In comparison, the model matching c3da main branches was rather similar with *k* = 0.15, *bf* = 0.1 and *radius* = 100*μm*.

The growth model by Baltruschat et al. (2020) was manually fitted to reproduce the main branches in the wild-type c3da neurons (**Figures 3A**). In order to do this the growth was first interrupted when the dendrite reached the number of main branch terminals in the real counterpart. The resulting dendritic total length served as a reference for finding good parameters. To account for synthetic morphologies grown in a given spanning area being systematically smaller than the original trees, the resulting model dendrites were slightly scaled to match the spanning area of their real counterparts.

Since the characteristic small terminal branches (STBs) of c3da dendrites were not well captured by the general growth model after resuming growth to math the total number of branches (**Figure 3B**), we implemented a transition to a second growth program after the main branches were grown. STBs were modelled by exploring which minimal changes needed to be introduced to the general growth model to obtain realistic total dendrite length, branch length distributions and distributions of STBs along the path from soma to the dendrite tip.

One viable model for the second growth step was found by restricting the reach of the targets to a close distance from the existing dendrite. This reach was inversely correlated with the local dendrite diameter *D* by 4.2*μm* – *D*. A stochasticity of the reach values was obtained by multiplying the reach by noise of 1*μm* ± 6*μm* low pass filtered with a Gaussian filter with a 60*μm* length constant. The reach was finally scaled by × 3.4 and capped at 10*μm* while reach values below 4.2*μm* were set to 0*μm* resulting in the characteristic STB-less stretches along c3da dendrites. It is important to note that we do not believe that the two growth steps happen subsequently but rather that their dynamics are intertwined. Furthermore, the second growth step had different parameters with *k* = 0.5, *bf* = 0.625 and without any further *radius* = *∞μm*. Most notably, the specific shape, angles and branch length distributions of STBs could only be reproduced when introducing a more fundamental change to the parameter *bf*. Here, instead of increasing cost with long paths to the dendrite root, the paths were measured in reference to the dendrite’s main branches resulting in mostly unbranched STBs directed towards the main dendrite (**Figure 3C**).

Mutant synthetic morphologies were grown using exactly the same two-step growth program as used for the wild-type morphologies. The only differences in morphology therefore come from the specific differences in dendrite spanning fields as well as from the number of main branches and total number of branches.

#### Data and Code Availability

The data and code in *Matlab* (www.mathworks.com) that support the findings in **Figures 3–6** of this study will be made available on publication.

## Supporting information

**Movie S1. Actin FRAP Recovery**

**A**, Representative time-lapse movie of FRAP recovery at the tip of a STB in a c3da neuron. Membrane mCherry signal in magenta and actin::GFP in green. Timeseries is 10 min with an image shown every 30 sec. The white circle indicates the area of bleachin and the white arrow the strong GFP signal at the tip of the growing branchlet. Scale bar is 5*μm*.

**Fig S1.**
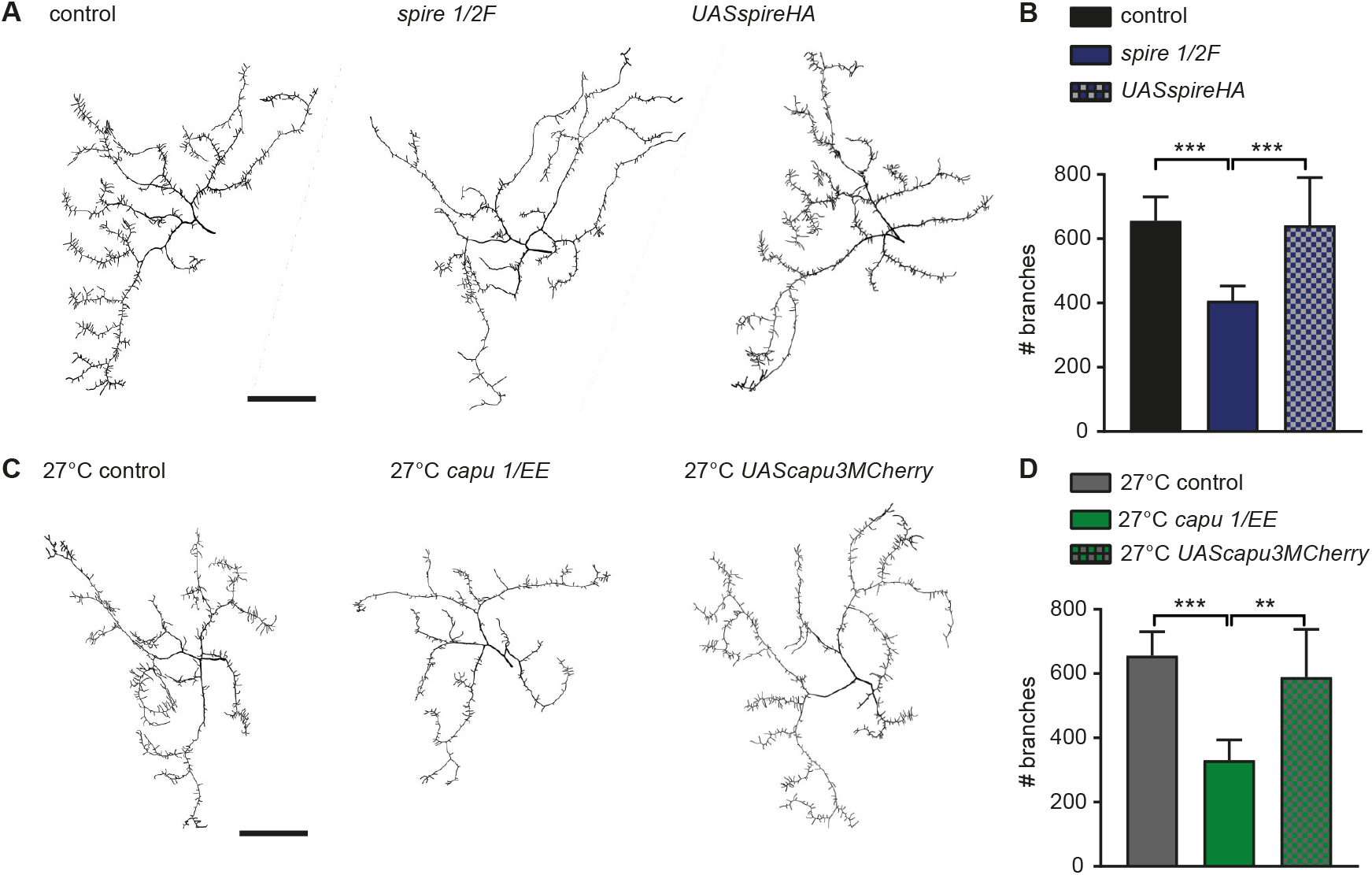
Spire and Capu Rescue. **A**, Representative tracings of control, *spire*^1^ /*spire*^2*F*^ mutant and *UASspirHA* rescue. **B**, Quantification of branch number. **C**, Representative tracings of control, *capu*^1^ /*capu^EE^* mutant and *UAScapu3MCherry* rescue. **D**, Quantification of branch number. (* is *p* < 0.05, ** is *p* < 0.01 and *** is *p* < 0.001). Scale bar is 100*μm*. *n* = 5 larva per genotype (see **Table 2** for genotypes).

**Fig S2.**
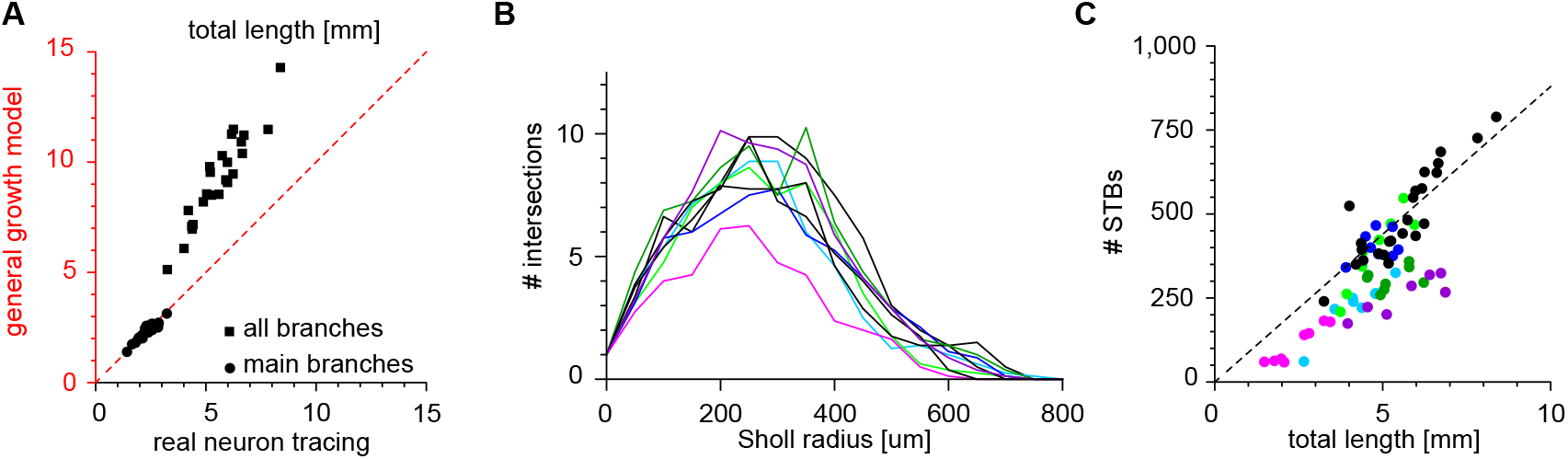
Further quantification of c3da neuron mutants. **A**, Direct comparison of total length in *mm* between reconstructions and c4da model. Dashed red line indicates same length. **B**, Sholl analysis of the main branches of control and mutant morphologies. **C**, The number of STBs against the total length for all controls and mutant tracings. Same colours as in **Figure 5**

**Fig S3.**
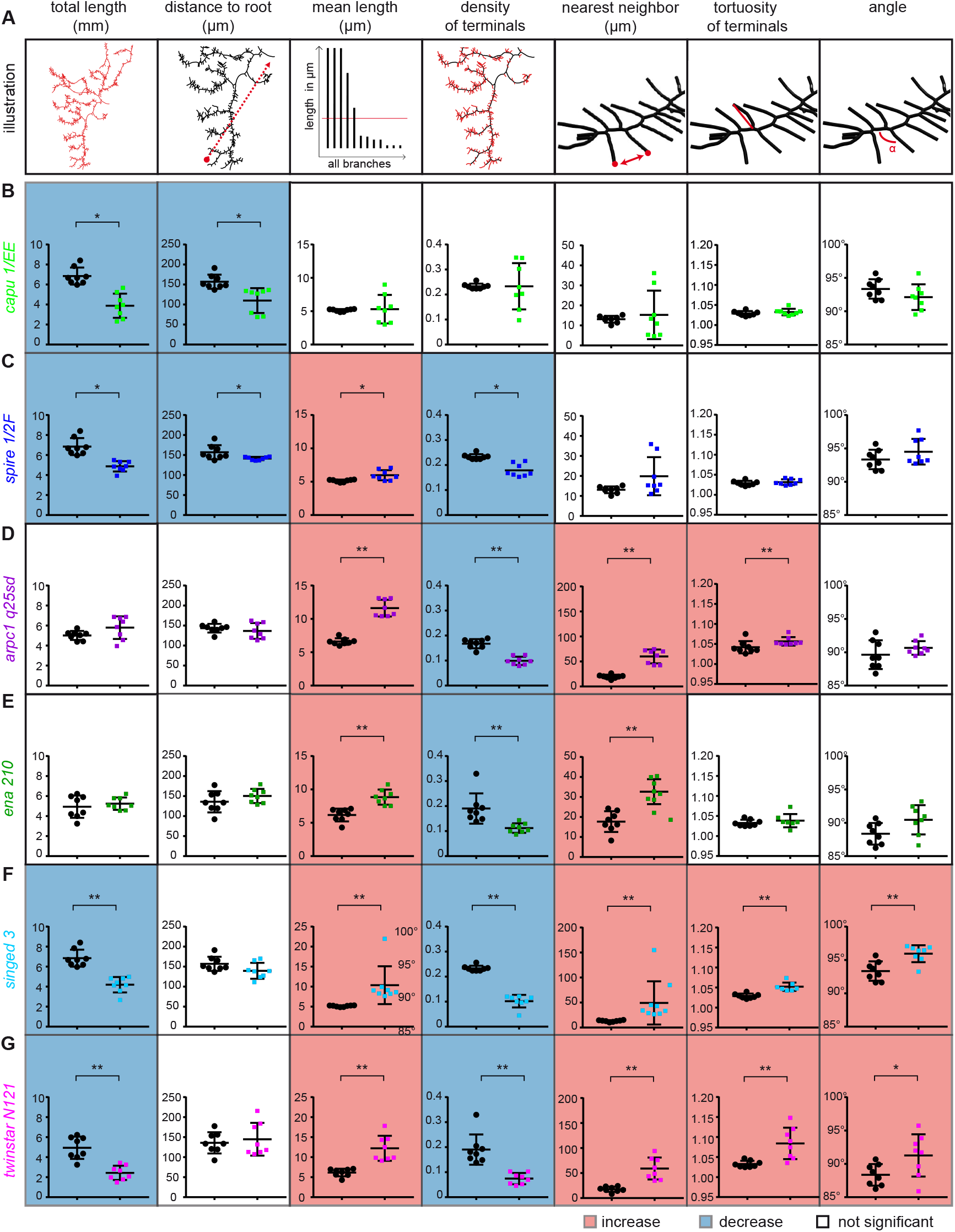
Morphological Analysis. **A**, Seven morphological measurements for the c3da neurons. **B-G**, The seven measurements for each ARP mutant compared to corresponding controls. (corrected *p* values * is *p* < 0.05, ** is *p* < 0.01 and *** is *p* < 0.001). The background is highlighted in blue for a significant decrease and in red for a significant increase.

**Fig S4.**
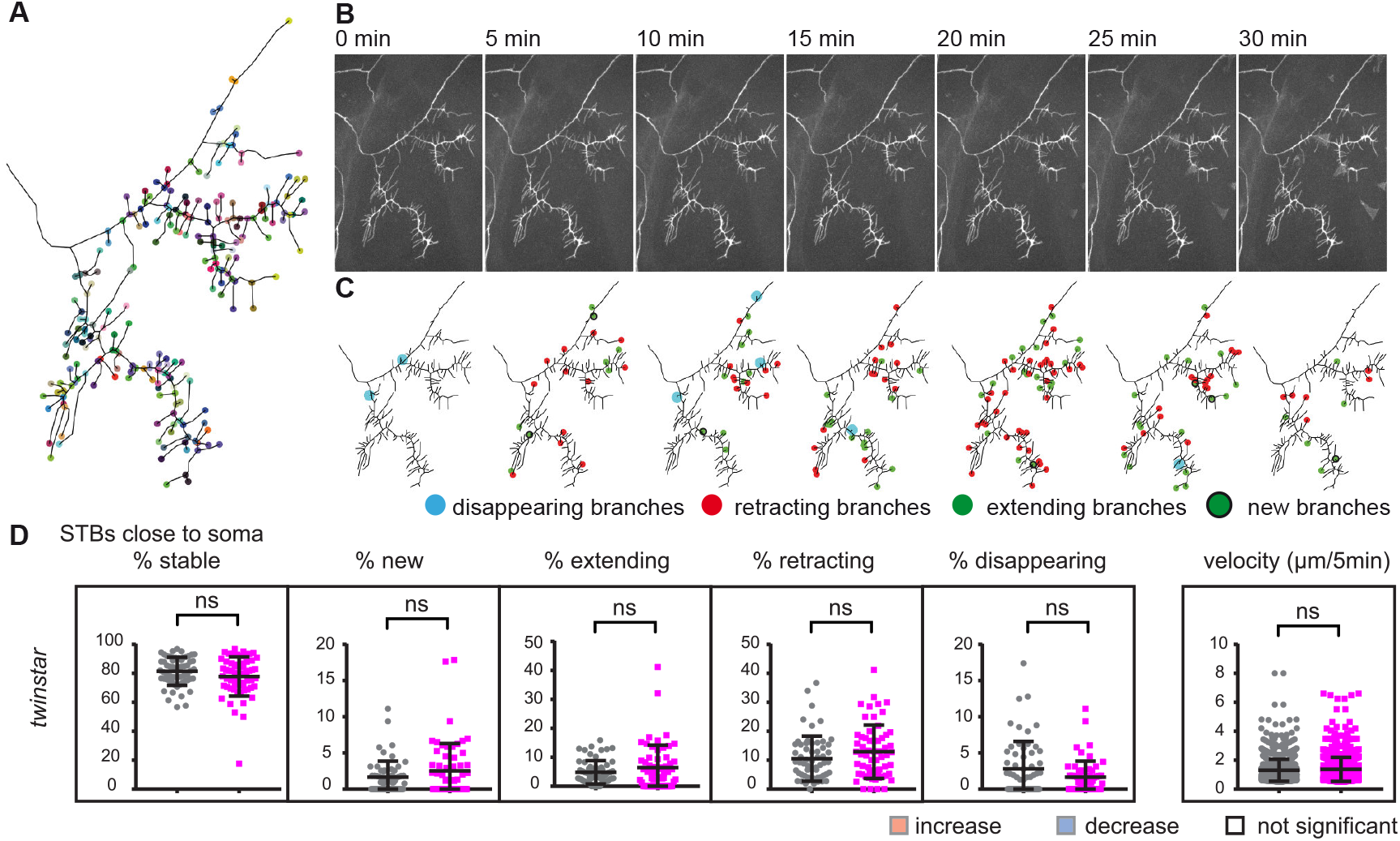
Time lapse analysis of control and specifically on few STBs close to the soma. **A**, Representative example of a tracing of a terminal region of a control c3da neuron. All branching points and terminal points are registered in the time-lapse series, illustrated as coloured points. **B**, Representative example of a control c3da neuron time-lapse series over 30*min* in 7 steps of 5*min* **C**, Tracing of the images in **B** with terminal branches that disappeared (blue), retracted (red) extended (green), or newly formed (green with black ring) from one time point to the next are marked with a dot in the corresponding colour (also shown in **Figure 6A**). **D**, Imaging and time-lapse analysis performed on the STBs close to the cell soma in *twinstar* mutants. Percentage of terminal branches that were stable, new, extending, retracting or disappearing within 30*min* of time-lapse for *twinstar* versus corresponding control (grey/black). Average velocity of a terminal branch, quantified as the average change in length (extension + retraction) in 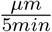 (see **Table 2** for genotypes).

